# Native molecule sequencing by nano-ID reveals synthesis and stability of RNA isoforms

**DOI:** 10.1101/601856

**Authors:** Kerstin C. Maier, Saskia Gressel, Patrick Cramer, Björn Schwalb

## Abstract

Eukaryotic genes often generate a variety of RNA isoforms that can lead to functionally distinct protein variants. The synthesis and stability of RNA isoforms is however poorly characterized. The reason for this is that current methods to quantify RNA metabolism use ‘short-read’ sequencing that cannot detect RNA isoforms. Here we present nanopore sequencing-based Isoform Dynamics (nano-ID), a method that detects newly synthesized RNA isoforms and monitors isoform metabolism. nano-ID combines metabolic RNA labeling, ‘long-read’ nanopore sequencing of native RNA molecules and machine learning. Application of nano-ID to the heat shock response in human cells reveals that many RNA isoforms change their synthesis rate, stability, and splicing pattern. nano-ID also shows that the metabolism of individual RNA isoforms differs strongly from that estimated for the combined RNA signal at a specific gene locus. And although combined RNA stability correlates with poly(A)-tail length, individual RNA isoforms can deviate significantly. nano-ID enables studies of RNA metabolism on the level of single RNA molecules and isoforms in different cell states and conditions.

## Main

In metazoan cells, a single gene locus can give rise to a variety of different RNA molecules that are generally referred to as isoforms. These RNA isoforms can differ in their 5’- and 3’-ends that arise from the use of different transcription start sites and polyadenylation sites, respectively ^1-4^. In addition, alternative splicing results in RNA isoforms that differ in the composition of their RNA body ^5,6^. Different mRNA isoforms can result in functionally different proteins. Vulnerabilities in splicing can lead to non-functional protein products. Diseases have been linked to alternative splicing, which can generate malignant RNA isoforms ^7^. Duchenne muscular dystrophy (DMD), for example, can be pinpointed to a single gene encoding the protein dystrophin. The underlying malignant RNA isoform exhibits a different splicing pattern and leads to a non-functional protein, which disrupts muscular cell integrity ^8^. Likewise, the three most common types of breast tumors are linked to exon skipping and intron retention ^9^.

RNA isoforms can also differ in their stability. The untranslated region of an RNA isoform can differ in length and contains regulatory elements ^10^. The length of the poly(A)-tail at the 3’-end of RNA isoforms can also differ and influence RNA stability ^11,12^, and this is relevant for disease as well ^13^. Finally, introns may be retained in RNAs and can influence stability ^14^.

Little is known however about the synthesis and stability of single RNA isoforms in cells. This is because the systematic characterization of RNA isoforms and their metabolism is technically difficult. In particular, the detection, quantification and estimation of the stability of RNA isoforms is essentially impossible with ‘short-read’ RNA sequencing methods because reads generally cannot be assigned to RNA isoforms. Also, alternative splicing patterns can be manifold and are difficult to identify using ‘short-read’ sequencing approaches ^15^. Finally, although the length of poly(A)-tails of RNAs can be measured genome-wide ^16,17^, they can currently not be obtained at the level of individual RNA isoforms.

The architecture of RNA isoforms has been addressed so far by ‘short-read’ RNA sequencing approaches such as DARTS ^18^, VastDB ^19^ and MPE-seq ^20^ to study alternative splicing or TIF-seq ^1,3^ to elucidate combinations of paired 5’- and 3’-ends of individual RNAs. More recent approaches include ‘long-read’ sequencing approaches on the PacBio SMRT Sequencing platform ^6^ or Oxford Nanopore Technologies nanopore sequencing platform ^5,21,22^. These methods however are not able to study the metabolism of individual RNA isoforms because they lack the ability to assign age to single reads.

Methods to measure the synthesis and stability of combined RNA for entire gene loci are available ^23-25^. Transient transcriptome sequencing (TT-seq) is a protocol that allows to distinguish newly synthesized from pre-existing RNA in human cells ^26^. TT-seq involves a brief exposure of cells to the nucleoside analogue 4-thiouridine (4sU). 4sU is incorporated into RNA during transcription, and the resulting 4sU-labeled RNA can be purified and sequenced to provide a snapshot of immediate transcription activity. This then enables to computationally infer RNA synthesis and stability at the level of the combined RNA signal from a gene locus.

Recent methods to assess RNA stability include SLAM-seq ^27^ and TimeLapse-seq ^28^. Like TT-seq, SLAM-seq and TimeLapse-seq involve an exposure of cells to 4sU for labeling of newly synthesized RNA. A chemical modification of the incorporated 4sU then allows for the identification of labeled RNA *in silico* without the need for purification. All of these methods, however, have limitations. First, sequencing reads can normally only be assigned to entire gene loci and not to RNA isoforms and thus only allow a combined RNA stability assessment. Second, they require template amplification, which can lead to an imbalance in measured sequences and information loss, e.g. modified RNA bases ^29^. Third, labeled RNA purification (TT-seq) and cDNA library preparation (TT-seq, SLAM-seq & TimeLapse-seq) can also introduce biases.

Therefore, monitoring RNA metabolism at the level of RNA isoforms requires a method that can detect individual RNA molecules. Recent advances in ‘long-read’ nanopore sequencing indeed enable the sequencing of single, full-length RNA molecules ^5^. Nanopore technology can directly sequence the original native RNA molecule with its modifications, may they be natural or acquired by metabolic RNA labeling. Moreover, the availability of the entire RNA and coding sequence (CDS) within a single read allows to unambiguously and directly determine exon usage ^30^. Direct RNA ‘long-read’ nanopore sequencing also has the potential to detect the position and length of the poly(A)-tail along with each single isoform.

Here we developed nanopore sequencing-based Isoform Dynamics (nano-ID), which combines metabolic RNA labeling with native RNA ‘long-read’ nanopore sequencing for RNA isoform detection. In combination with computational modeling and machine learning this allows for a full characterization of RNA isoforms dynamics. nano-ID can identify and quantify RNA isoforms along with their synthesis rate, stability and poly(A)-tail length in the human myelogenous leukemia cell line K562. We show that this is possible with nano-ID in a quantitative manner in steady state and also during the transcriptional response to heat shock. nano-ID is able to resolve the dynamic metabolism of RNA isoforms upon heat shock and demonstrates the need for individual RNA isoform assessment. Taken together, nano-ID can be used to elucidate a largely unexplored complex layer of gene regulation at the level of single native RNA isoforms and their metabolism.

## Results

### Experimental design

To monitor the metabolism of RNAs at the level of single isoforms, we sought to combine metabolic RNA labeling with direct, single-molecule RNA nanopore sequencing (**Figure 1a**). By culturing cells in the presence of a nucleoside analogue, cells will take up and incorporate the analogue in nascent RNA during transcription, allowing to distinguish newly synthesized RNA isoforms from pre-existing RNA isoforms *in silico* based on the quantification of analogue-containing subpopulations. This will allow to infer the synthesis rate and stability of single RNA isoforms. In order to dynamically characterize functional and fully processed RNA transcripts, we decided to measure poly-adenylated RNA species. The library preparation kit offered by Oxford Nanopore Technologies for direct RNA sequencing (SQK-RNA001) is specifically optimized for this purpose. A 3’ poly(A)-tail specific adapter is ligated to the transcript in a first step. Then a second sequencing adapter equipped with a motor protein is ligated to the first adapter. The preparation of RNA libraries from biological samples for direct RNA nanopore sequencing is established and can be carried out within 2h ^31^. Major challenges that we faced were however the search of a suited nucleoside analogue for RNA labeling and the detection of labeled RNA isoforms, provided that the labeling efficiency is known to be limited to about 2-3%, i.e. only two or three out of 100 natural nucleotides are replaced by the analogue ^32^.

**Figure 1.**
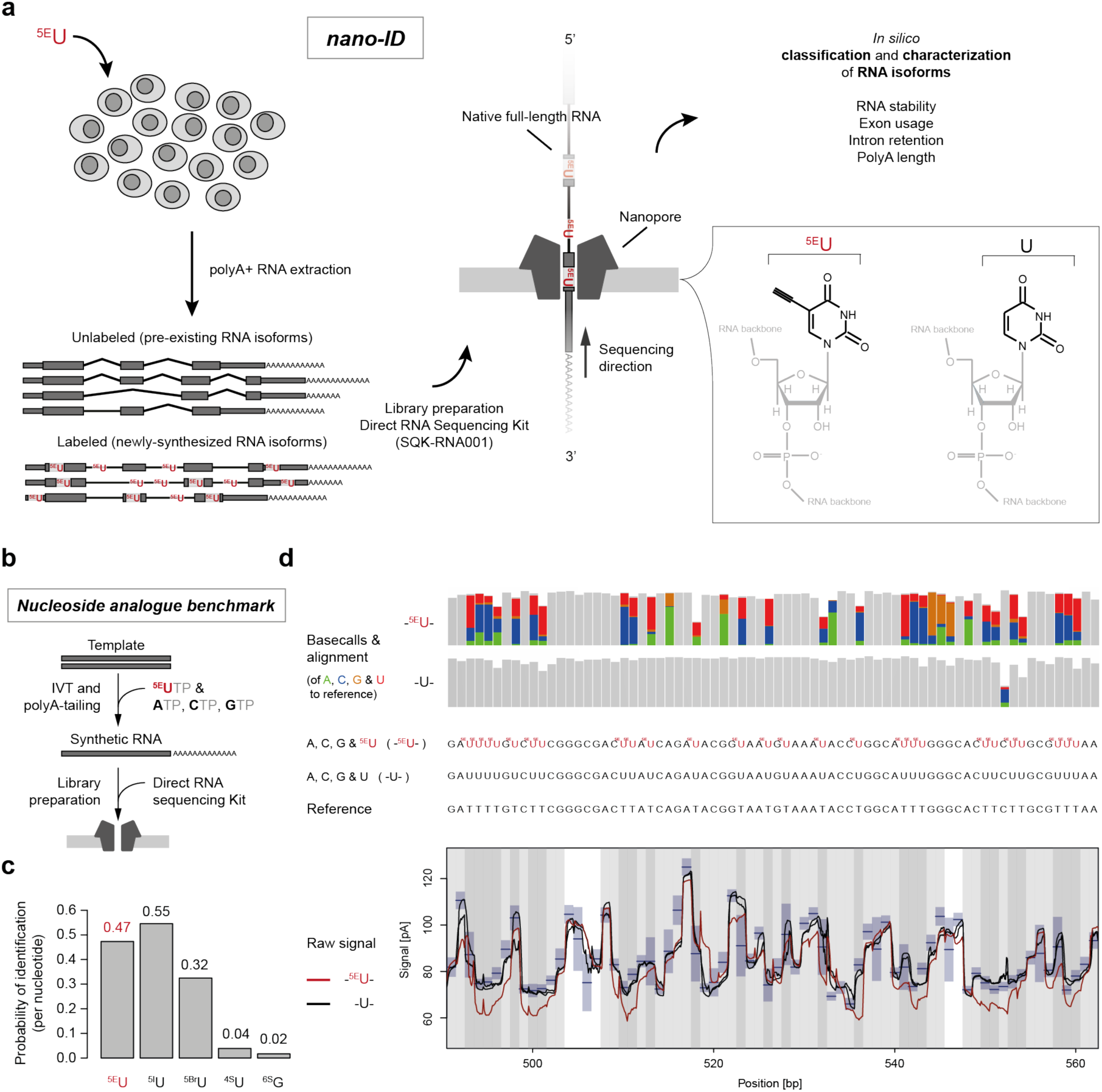
Nanopore sequencing-based Isoform Dynamics (nano-ID) combines metabolic RNA labeling with ‘long-read’ nanopore sequencing of native RNA molecules. (a) Experimental schematic of ^5E^U-labeled RNA isoforms subjected to direct RNA ‘long-read’ nanopore sequencing. Metabolic labeling of human K562 cells with the nucleoside analogue 5-Ethynyluridine (^5E^U) *in vivo*. Newly-synthesized RNA isoforms will incorporate ^5E^U instead of standard uridine (U) residues. This allows to distinguish the newly synthesized RNA isoforms (Labeled) from pre-existing RNA isoforms (Unlabeled) *in silico* after sequencing the native full-length molecules on an array of nanopores ^5^. ^5E^U containing RNA isoforms are computationally traceable and thus allow classification. Identification and quantification of RNA isoforms subsequently enable assessment of RNA stability, exon usage, intron retention and polyA-tail length. (b) Experimental schematic to derive synthetic RNAs for nucleoside analogue benchmark. RNAs were *in vitro* transcribed using either the standard bases A, U, C, G as a control, or one of the natural bases was exchanged for a nucleoside analogue (shown for ^5E^U). (c) Barplot showing the probability of nucleoside analogue identification compared to natural UTP/GTP based on base-miscalls (**Methods**) of all tested nucleoside analogues (^5E^U, 5-bromouridine (^5Br^U), 5-iodouridine (^5I^U), 4-thiouridine (^4S^U) and 6-thioguanine (^6S^G)). (d) Upper panel: Base miscalls (colored vertical bars) of the standard base-calling algorithm for synthetic RNAs containing ^5E^U instead of U (-^5E^U-, 3.563 molecules) and synthetic control RNAs (-U-, 15.840 molecules) in comparison to the original sequence (Reference) of an exemplary region on synthetic RNA ‘Spike-in 8’ (**Methods, Supplementary Table 3**). Middle panel: Synthetic RNA sequences with (-^5E^U-) and without ^5E^U (-U-) depicted above the reference sequence (Reference). Lower panel: Alignment of the raw signal readout of the nanopore in pico-Ampere [pA] to the reference sequence. Synthetic control RNAs (-U-) are shown in black. ^5E^U containing synthetic RNAs are shown in red (-^5E^U-). ^5E^U containing synthetic RNAs show a clear deviation from the expected signal level in blue. Blue boxes indicate mean and standard deviation of the pore model on which the original base-calling algorithm is based.

### 5-Ethynyluridine (^5E^U) can be detected in RNA by nanopore sequencing

To investigate if nucleoside analogues incorporated into RNA are detectable in the nanopore, we used synthetic RNAs derived from the ERCC RNA spike-in mix (Life Technologies). These synthetic RNAs of an approximate length of 1,000 nucleotides were chosen with similar U content (**Supplementary Table 3**). RNAs were transcribed *in vitro* using either the standard bases A, U, C, G as a control, or with one of the natural bases exchanged for a nucleoside analogue (**Figure 1b**, **Methods**). Subsequently, we subjected these synthetic RNAs to direct RNA nanopore sequencing (**Supplementary Figure 1a-b**). We compared the nucleoside analogues 5-Ethynyluridine (^5E^U), 5-bromouridine (^5Br^U), 5-iodouridine (^5I^U), 4-thiouridine (^4s^U) and 6-thioguanine (^6s^G). To this end we used the base-called and mapped direct RNA sequencing results to calculate how probable the identification would be on the level of single nucleotides. In particular, we compared the error rate in single nucleotide base-calls of nucleoside analogues to that of natural U or G (**Figure 1c, Methods**).

The thiol-based analogues, ^4s^U and ^6s^G, showed lower incorporation efficiencies during *in vitro* transcription (IVT) and led to blockages during nanopore sequencing. ^5E^U and ^5I^U could be detected to a similar extent by nanopore sequencing, whereas ^5Br^U was less easily recognized (**Figure 1c**). Since ^5E^U is not toxic to cells ^32-34^, we used ^5E^U for a more detailed analysis. Approximately 50% of all U positions in ^5E^U-containing synthetic RNAs are consistently miscalled by the standard base-calling algorithm and can thus be discerned from U (**Figure 1d, Supplementary Figure 1b**). This is clearly visible in the raw data. Aberrations caused by stretches of RNA containing ^5E^U are distinguishable from stretches of RNA containing the naturally occurring U in the nanopore (**Figure 1d**). Taken together, ^5E^U-based RNA labeling is well suited for nanopore sequencing.

### Detection and sequencing of newly synthesized RNA isoforms

We next investigated whether it is possible to use metabolic RNA labeling with ^5E^U in human cells to detect single RNA molecules by nanopore sequencing. Calculations on the direct RNA nanopore sequencing results of the ^5E^U-containing synthetic RNAs showed that RNAs are recognizable as ^5E^U containing with a probability of 0.9 when a minimum length of 500 nucleotides is reached (**Supplementary Figure 1c-d**). This covers the vast majority (93%) of all mature RNAs in the human organism (UCSC RefSeq GRCh38).

We then established direct RNA nanopore sequencing in the human myelogenous leukemia cell line K562. We cultured K562 cells in the presence of ^5E^U for 60 minutes (^5E^U 60 min) in 4 biological replicates (**Methods**). For comparison, we created 3 biological replicates exposed to ^5E^U labeling for 24 h (^5E^U 24 h) and 3 biological replicates that were not labeled (Control). After standard base-calling, we could map reads to support 13,110 RefSeq annotated transcription units (RefSeq-TUs, **Methods**), 8,098 of these were supported in all conditions and 1,726 were supported in all samples.

All combined samples were then used to perform a full-length alternative RNA isoform analysis by means of the FLAIR algorithm ^22^. This allows defining instances of unique exon-intron architecture with unique start and end sites in human K562 cells. Raw human direct RNA nanopore reads were corrected with the use of short-read sequencing data (RNA-seq) to increase splice site accuracy. We could detect 33,199 distinct RNA isoforms with an average of 3 isoforms per gene. This shows that direct RNA nanopore sequencing uncovers individual RNA isoforms in human K562 cells (**Figure 2**) with high reproducibility (**Supplementary Figure 2**).

**Figure 2.**
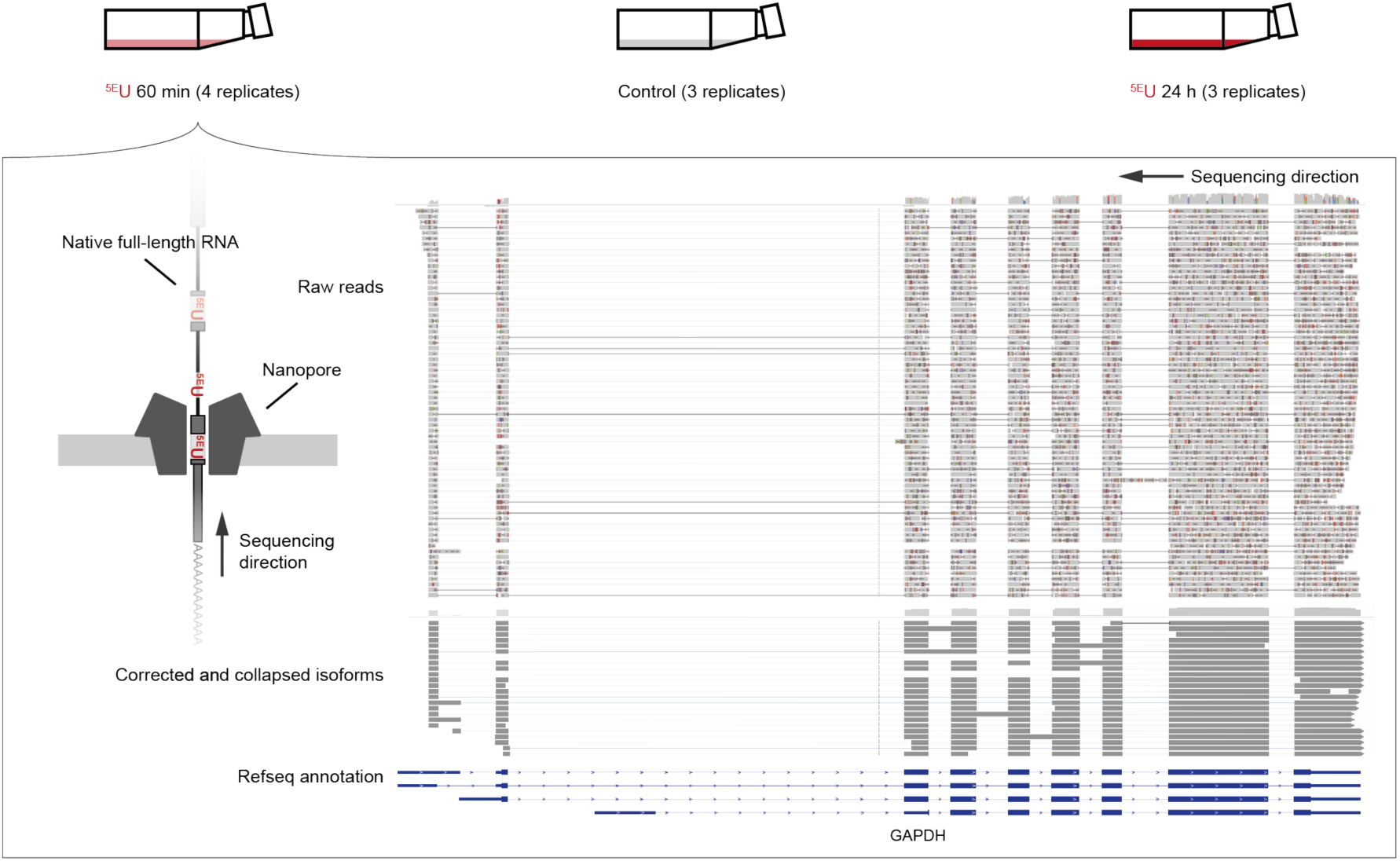
Direct RNA ‘long-read’ nanopore sequencing of ^5E^U-labeled RNA isoforms in human K562 cells. Upper panel: Illustration of the experimental set-up. Human K562 cells were cultured in the presence of the nucleoside analogue ^5E^U for 60 minutes (^5E^U 60 min, 4 replicates) and 24 h (^5E^U 24 h, 3 replicates). Control samples were not labeled (Control, 3 replicates). Lower panel: Genome browser view of direct RNA ‘long-read’ nanopore sequencing results of the human GAPDH gene locus on chromosome 12 (∼8 kbp, chr12: 6,532,405-6,540,375) visualized with the Integrative Genomics Viewer (IGV, version 2.4.10; human hg38) **^52^**. From top to bottom: raw nanopore sequencing reads (light grey, shown are typical aligned raw reads below the accumulated coverage of all measured reads), corrected and collapsed isoforms (dark grey) determined with the FLAIR algorithm ^22^ based on raw reads and RefSeq GRCh38 annotation (blue).

### A neural network identifies newly synthesized RNA isoforms

The next step was to derive a computational method that could classify each sequenced RNA molecule into one of two groups, newly synthesized (^5E^U-labeled) or pre-existing (unlabeled) RNA. To this end, the nucleoside analogue ^5E^U had to be detected in RNA molecules. This would allow the quantification of RNA isoforms generated during the ^5E^U labeling pulse. Due to the high error rate of nanopore sequencing, a single ^5E^U base-call is inappropriate as an indicator. We rather used the raw signal of the entire RNA nanopore read, including the base-calls and the alignment, to discriminate labeled from unlabeled RNAs. This discrimination was implemented as a classifying neural network. We developed a custom multi-layered data collection scheme to train a neural network for the classification of human RNA isoforms under the assumption that the ^5E^U 24 h samples solely contain labeled reads and the fact that the Control samples solely contain unlabeled reads (**Figure 3a, Methods**).

**Figure 3.**
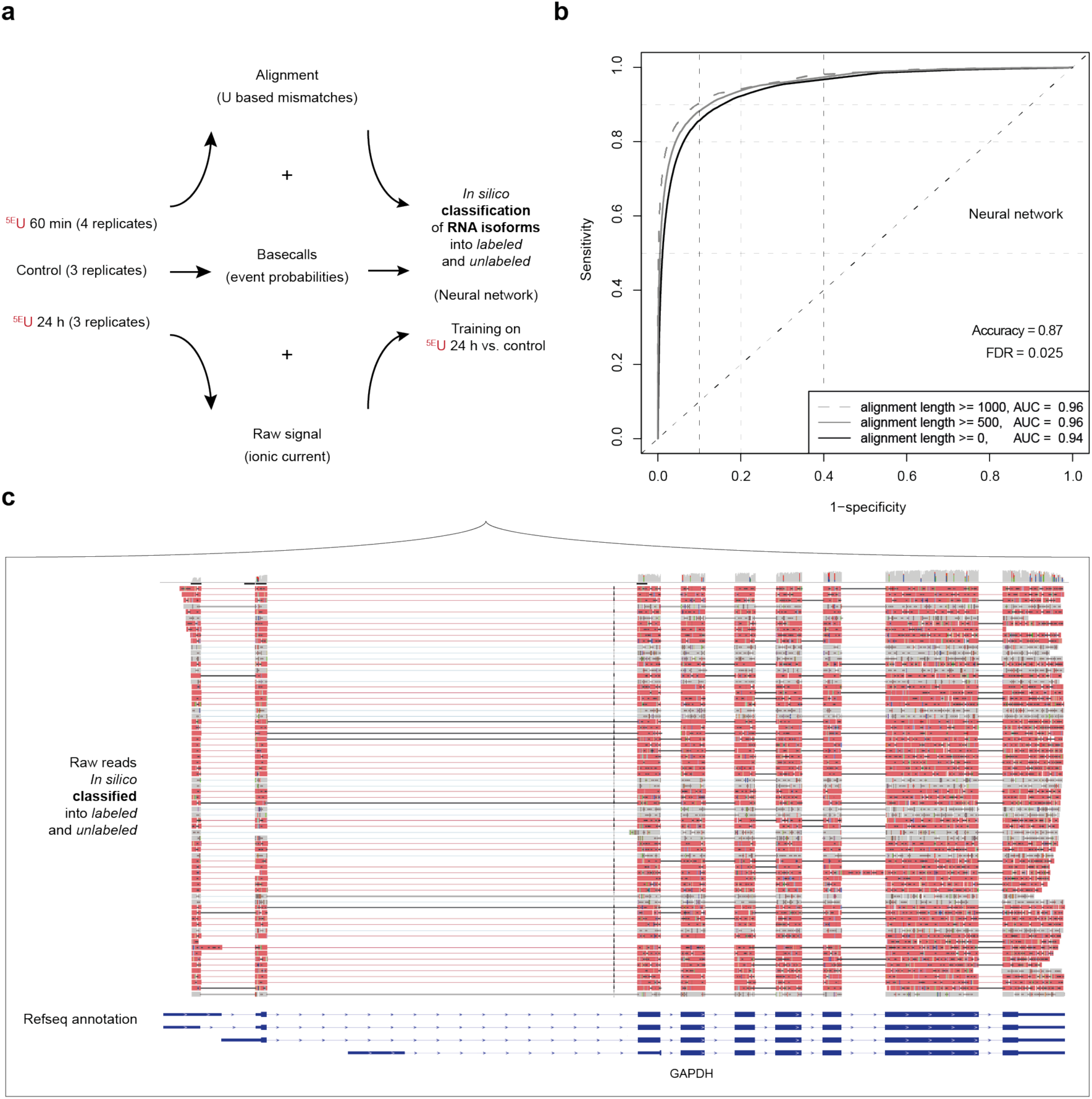
Neural network based classification of human RNA isoforms into ^5E^U-labeled and unlabeled. (a) Multi-layered data collection scheme. Parameter collection of human K562 samples (^5E^U 60 min, Control & ^5E^U 24 h) was realized on three different layers: Raw signal (ionic current), base-call event probabilities and alignment derived U based mismatch properties (**Methods**). Neural network was trained on the ^5E^U 24 h versus Control samples with an accuracy of 0.87 and a false discovery rate (FDR) of 0.025 and used to classify reads of the ^5E^U 60 min samples into ^5E^U-labeled and unlabeled. (b) ROC analysis of 5-fold cross-validated neural network training. Plot shows ROC curves (1 – specificity versus sensitivity) for all reads of the test set (black, alignment length >=0 nt, AUC = 0.94), for reads with an alignment length larger than 500 nt (grey, alignment length >=500 nt, AUC = 0.96) and for reads with an alignment length larger than 1000 nt (dashed grey, alignment length >=1000 nt, AUC = 0.96). (c) Genome browser view of classified direct RNA ‘long-read’ nanopore sequencing reads of the human GAPDH gene locus on chromosome 12 (∼8 kbp, chr12: 6,532,405-6,540,375) visualized with the Integrative Genomics Viewer (IGV, version 2.4.10; human hg38) **^52^**. Unlabeled reads are shown in grey, ^5E^U-labeled reads are shown in red.

We then trained a neural network (**Methods**) on the ^5E^U 24 h versus Control samples with an accuracy of 0.87 and a false discovery rate (FDR) of 0.025 (5-fold cross-validated). A ROC analysis (1 – specificity versus sensitivity) for all reads of the test set showed an area under the curve (AUC) of 0.94. For reads with an alignment length larger than 500 nt and 1,000 nt the AUC improved to 0.96 (**Figure 3b, Supplementary Figure 3a, b**). Subsequently we used the trained neural network to classify reads of the ^5E^U 60 min samples into ^5E^U-labeled and unlabeled. Taken together, ^5E^U containing RNA isoforms are computationally detectable with high accuracy (**Figure 3c**). For validation purposes, we used another machine learning approach. We trained a random forest on the same data, which yielded similar results (**Supplementary Figure 3c, d**). Thus, we were able to determine for each single RNA molecule if it has been produced during ^5E^U labeling or before, with a low false discovery rate (**Figure 3c**).

### nano-ID provides the stability and poly(A) tail length of RNA isoforms

The ability to distinguish newly synthesized and pre-existing RNA molecules allowed us to derive estimates for the stability of RNA isoforms. For each single direct RNA nanopore read we were able to assign the RNA isoform it reflects. Additionally, we were able to assess the stability of RNA for single RNA isoforms by applying a first-order kinetic model (**Methods, Supplementary Figure 3e-f**) to derive estimates for RNA isoform specific synthesis and stability. This can be done based on the number of reads classified as ^5E^U-labeled and unlabeled by the neural network. Taken together, nano-ID has the capability to infer synthesis and stability of individual RNA isoforms in different cell states and conditions, and thus to monitor their dynamic metabolism.

Moreover, we developed an algorithm to determine poly(A)-tail lengths for each RNA isoform (**Figure 4**). This is possible by estimating the dwell time of the poly(A)-tail in the nanopore by factoring in the measurement frequency in kHz and the speed of RNA translocation through the nanopore (**Methods**). Sequencing adaptor ligation in the direct RNA nanopore sequencing library preparation guarantees full-length poly(A)-tails because ligation of the adapter would not be successful otherwise. The resulting poly(A)-tail length distribution is in line with the current literature ^16^ (**Figure 4a**) and reveals a pattern that likely corresponds to the 26 nucleotide footprint of the poly(A) binding protein (**Supplementary Figure 4a**) ^35^. The direct RNA nanopore sequencing kit contains the so-called RNA calibration strand (RCS). The RCS is a synthetic RNA with a poly(A)-tail of exactly 30 adenines. Using the RCS of the direct RNA nanopore sequencing kit, we could assess the accuracy of the poly(A)-tail length estimates (coefficient of variation 0.63). Our algorithm derives this length for the added RCS subpopulation (**Figure 4b**). Taken together, nano-ID reveals the synthesis, stability, and poly(A) tail length for individual RNA isoforms in human cells.

**Figure 4.**
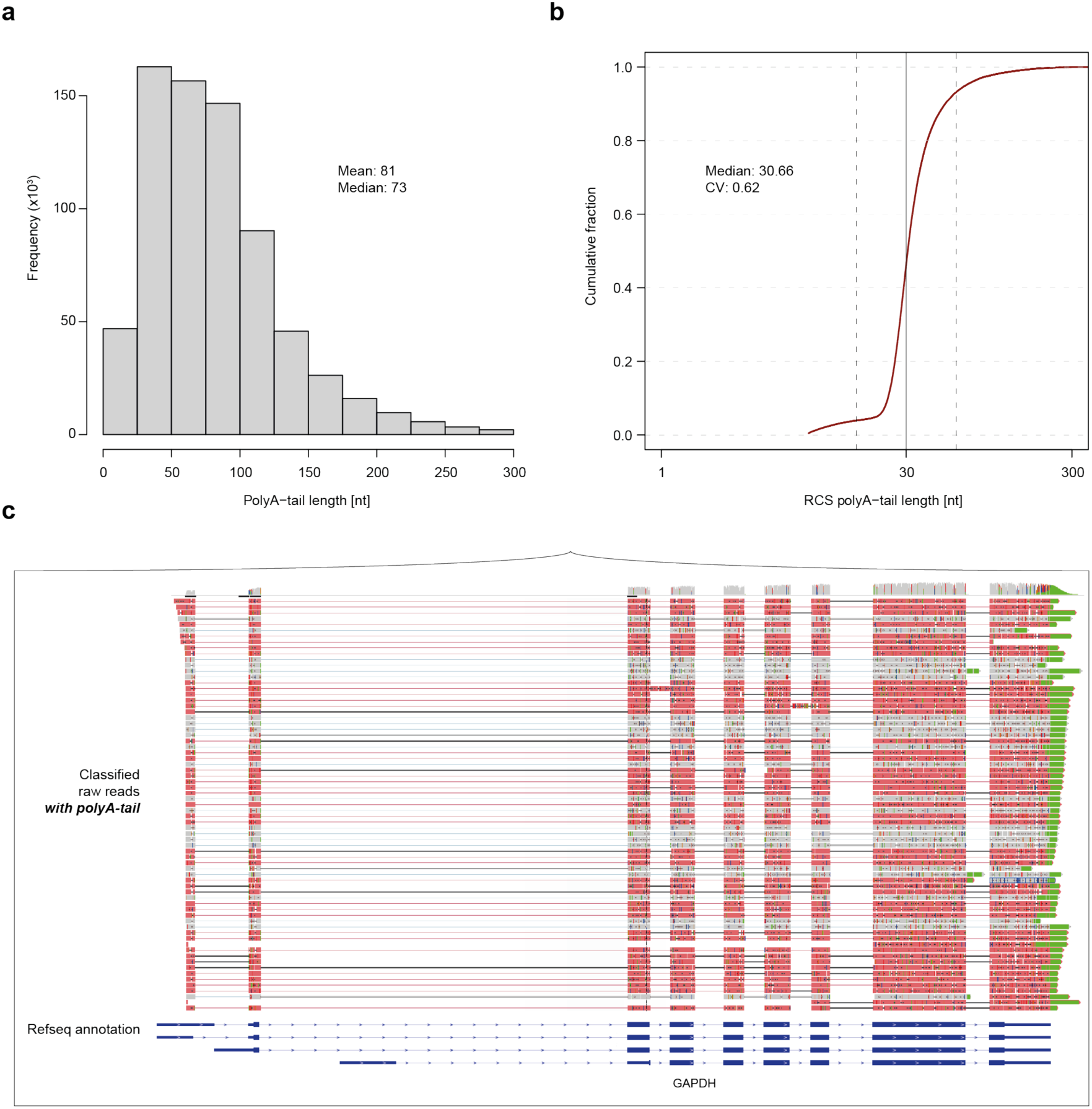
Poly(A)-tail length determination of human RNA isoforms. (a) Histogram of poly(A)-tail length estimates of 714,536 RNA isoforms (mean: 81 nt, median: 73 nt). (b) Cumulative distribution function of poly(A)-tail length estimates of the RNA calibration strand (RCS, yeast derived spike-in RNAs that are equipped with a poly(A)-tail of exactly 30 adenines (ONT, SQK-RNA001)). Vertical solid black line indicates optimal result of 30 nt (median: 30.6, coefficient of variation: 0.62). Vertical dashed black lines indicate 2-fold in either direction. (c) Genome browser view of classified direct RNA ‘long-read’ nanopore sequencing reads with poly(A)-tail (green) of the human GAPDH gene locus on chromosome 12 (∼8 kbp, chr12: 6,532,405-6,540,375) visualized with the Integrative Genomics Viewer (IGV, version 2.4.10; human hg38) **^52^**.

### nano-ID monitors RNA isoform dynamics during heat shock

To demonstrate the advantages of nano-ID, we subjected human K562 cells to heat shock (42 °C) for 60 min in the presence of ^5E^U (^5E^U 60 min HS) (**Figure 5a**). The heat shock response provides a well-established model system ^36-41^ (**Supplementary Figure 5**). We first asked whether RNA isoforms do retain more introns after heat shock as this was shown in the mouse system ^42^. Indeed, we observed widespread intron retention which significantly increased upon heat shock (**Figure 5b**). Although intron retention generally influences the stability of an RNA, it does not explain changes in RNA isoform stability upon heat shock (**Figure 5c**). This finding is consistent with the idea that specific RNA elements occurring only in specific RNA isoforms influence RNA stability.

**Figure 5.**
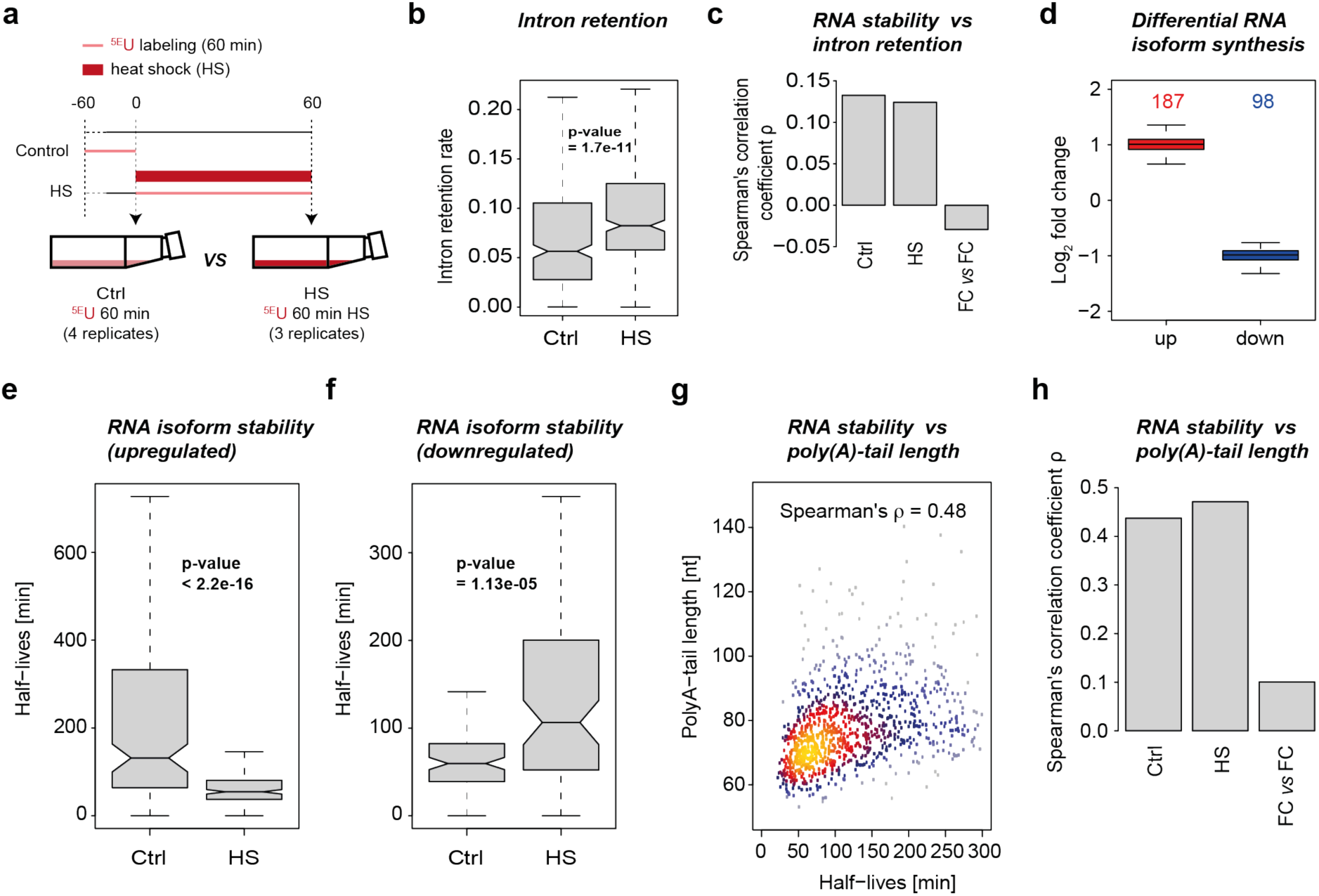
nano-ID monitors RNA isoform dynamics during heat shock. (a) Experimental set-up of the heat shock treatment (60 min at 42 °C) in human K562 cells. (b) Boxplot shows intron retention rate (**Methods**, min 5% in either condition) of 358 gene loci comparing heat shock (^5E^U 60 min HS) against control (^5E^U 60 min). (c) Bar plot shows correlation (Spearman’s rank correlation coefficient) of RNA half-lives and intron retention ratios before and after heat shock (1,027 loci). The third bar shows the correlation of their respective folds. (d) Boxplot shows upregulated (red) and downregulated (blue) RNA isoforms upon 60 min of heat shock (42 °C). A minimum fold change of 1.25 and a maximum p-value of 0.1 was set for calling a significant expression change. (e) Boxplot shows half-lives of significantly upregulated RNA isoforms comparing heat shock (^5E^U 60 min HS) against the control (^5E^U 60 min). (f) As in (e) for significantly downregulated RNA isoforms. (g) Scatter plot with color-coded density of RNA half-lives and RNA poly(A)-tail lengths in both conditions. Shown are 1,230 highly expressed RefSeq GRCh38 annotated genes. Correlation is calculated as Spearman’s rank correlation coefficient (0.48) rounded to the second decimal. (h) As in (c) using the RNA poly(A)-tail lengths (1,230 loci).

We next asked if RNA isoform synthesis is altered by heat shock and observed significant differential RNA isoform synthesis for 285 isoforms (fold change > 1.25 and p-value < 0.1). 187 RNA isoforms were significantly upregulated, while 98 were downregulated (**Figure 5d**). RNA isoforms that changed their synthesis during heat shock were also observed to alter their stability (**Figure 5e-f**). In particular, RNA isoforms that were upregulated in their synthesis during heat shock also showed a lower stability, and the other way around, resembling typical stress response behavior ^24^. The destabilization of upregulated RNA isoforms is likely to ensure their rapid removal toward the end of the stress response. Similarly, downregulated RNA isoforms are stabilized, perhaps to preserve them for translation at later stages.

### nano-ID reveals the biogenesis of RNA isoforms

Although standard native RNA isoform sequencing can reveal isoforms present in a sample after perturbation, it cannot distinguish whether these isoforms were derived by synthesis, stability, splicing, or any combination of these. nano-ID however is able to disentangle these parameters. For example, although we observe a general increase in intron retention upon heat shock, we find exceptions at the level of RNA isoforms. This can be clearly seen at the human C1orf63 gene locus (**Supplementary Figure 5g**). Here, the majority of reads, that retain the entire 3^rd^ intron, were newly synthesized in the control samples. It is however unclear if this intron will be retained throughout the existence of these RNA molecules. Investigation of the same gene locus upon heat shock showed that the vast majority of reads were pre-existing RNAs. This indicates that this RNA is not transcribed anymore upon heat shock and allows for the conclusion that intron retention is not altered, rather, less introns are seen retained when only old RNA is detected. Taken together, this shows that nano-ID is able to resolve the dynamic behavior of RNA isoforms upon stimuli that could not be seen otherwise. It demonstrates the need for individual RNA isoform detection and classification into newly synthesized and pre-existing molecules. By providing information on the age of RNA molecules, nano-ID enables an analysis of the biogenesis of RNA isoforms.

### The metabolism of individual RNA isoforms differs from combined RNA estimates

To demonstrate the importance of individual RNA isoform assessment, we first derived estimates for the half-lives of combined RNAs that stem from entire gene loci under steady state conditions (**Methods, Supplementary Figure 3e-f**). We found a robust correlation of combined RNA stability with poly(A)-tail length (Spearman’s rank correlation coefficient 0.48) (**Figure 5g**). We now asked whether changes in RNA stability would also be reflected in changes in poly(A)-tail length upon heat shock, and this was not the case (**Figure 5h**). Instead, we found genes that showed the opposite behavior to the overall correlation as demonstrated for the human HSPB1 locus (**Figure 6a-b**). Here, destabilization of combined RNAs is accompanied by lengthening of the poly(A)-tail. This view changes dramatically when considering individual RNA isoforms (**Figure 6c**). For those three RNA isoforms at the human HSPB1 gene locus for which stability estimates were supported by all 3 biological replicates (**Methods**) we found that poly(A)-tails were generally longer. RNA stability however was decreased for 2 out of the 3 RNA isoforms and increased for the third. This clearly indicates the need for detailed individual RNA isoform assessment as individual RNA isoforms can lead to functionally distinct protein variants. Thus, it is crucial to also study the behavior of individual RNA isoforms instead of breaking it down to the combined view of the entire gene locus.

**Figure 6.**
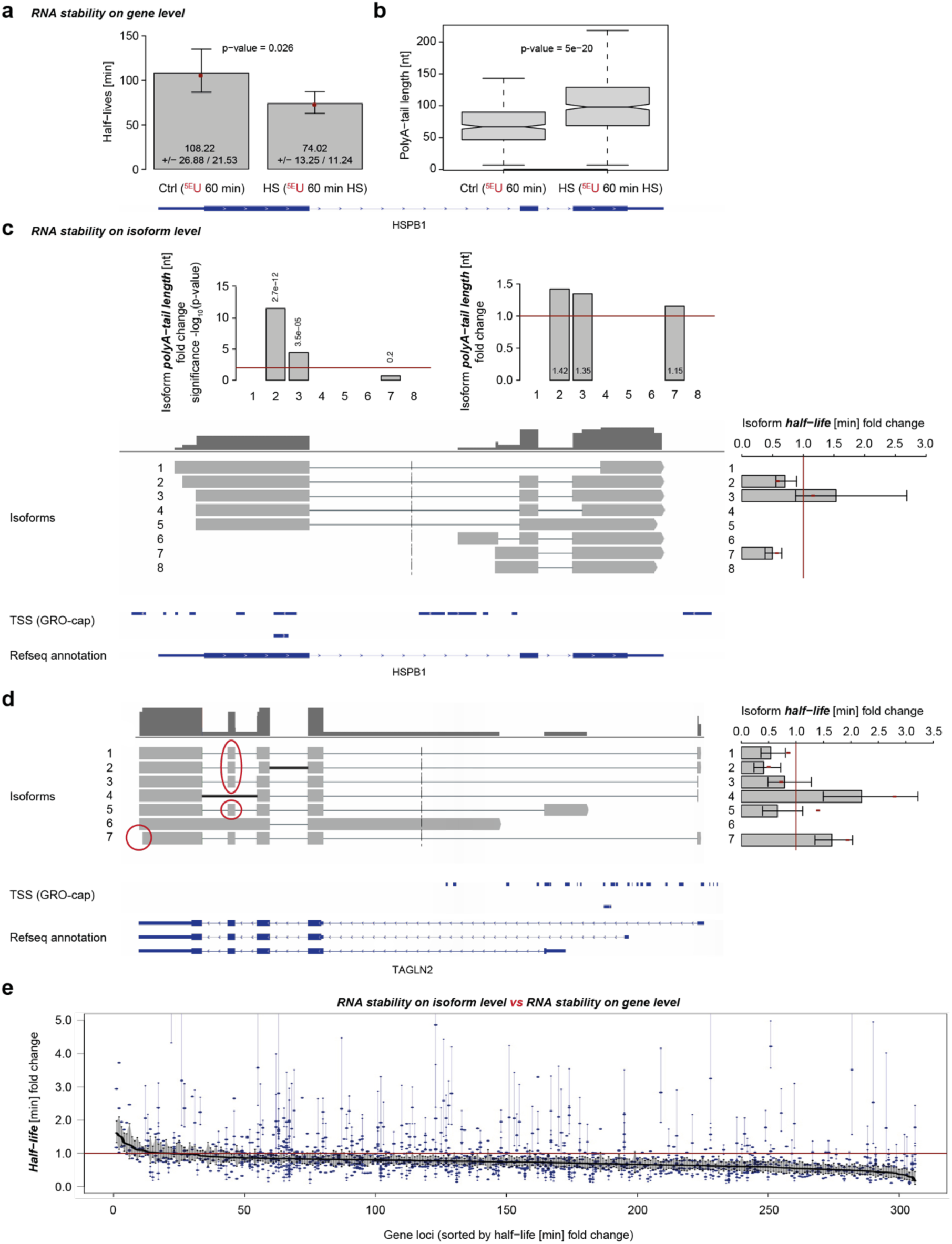
nano-ID resolves the characteristics of individual RNA isoforms. (a) Boxplot shows half-life estimates of RNAs from the human HSPB1 gene locus (chr6:31,813,514-31,819,942) comparing heat shock (HS, ^5E^U 60 min HS) against control (Ctrl, ^5E^U 60 min). Standard deviation is shown as error bars. Red points depict half-life estimate of merged replicates in each condition. (b) Boxplot shows the poly(A)-tail length distributions of RNAs from the human HSPB1 gene locus. 437 RNAs from heat shocked samples (HS, ^5E^U 60 min HS) are compared to 341 RNAs in the respective control sample (Ctrl, ^5E^U 60 min). (c) Schematic shows direct RNA nanopore sequencing derived RNA isoforms at the human HSPB1 gene locus above annotated transcription start sites (TSSs) from published GRO-cap data generated in K562 cells **^2^** and RefSeq GRCh38 annotation. Bar plots show RNA isoform half-life fold changes, poly(A)-tail length fold changes and their respective significance as standard deviation (error bars) or -log_10_(p-value). Red lines indicate no fold change or -log_10_(p-value) with p-value 0.01. (d) As in (c) for RNA isoforms at the human TAGLN2 gene locus (chr1:159,916,107-159,927,542). (e) Half-life fold change (y-axis) depicted for RNAs encoded by 306 high confident gene loci (x-axis). All estimates are supported across biological replicates (n≥3) and conditions (n=2). Half-life estimates for RNA encoded by the entire gene loci (combined) are depicted as a black line (sorted in decreasing order). Blue dots represent individual RNA isoform half-life estimates at respective gene loci (1,169 isoforms in total). Perpendicular blue and black lines represent standard deviations of individual estimates. For individual RNA isoform half-life estimates, standard deviations are only shown if not overlapping with the standard deviation of the respective combined half-life estimates (black).

As a second example, we picked RNA isoforms at the human TAGLN2 gene locus (**Figure 6d**). We could identify 7 different RNA isoforms and reliably calculate RNA stability for 6 RNA isoforms. Two of them were stabilized upon heat shock, 4 of them were destabilized. All 4 destabilized RNA isoforms include the second to last exon, which might cause this change in stability. RNA isoform 7 is an exception to this observation as it is stabilized upon heat shock. It, however, also contains a 3’ UTR that is 42 bases shorter than all the other RNA isoforms. We asked whether there is differential behavior of individual RNA isoforms genome-wide or if RNA isoforms generally reflect the changes in stability of the combined RNA from their respective gene loci. To that end, we compared RNA stability estimates of individual RNA isoforms to those from combined RNAs and found that the dynamics of individual RNA isoform during heat shock varies globally (**Figure 6e**, **Supplementary Figure 6**). Taken together, this shows that conclusions can be misleading when combined RNAs are used and how much can be learned on the level of single RNA isoforms by using nano-ID.

## Discussion

Here we develop nano-ID, a method that allows for dynamic characterization of functional and fully processed RNA isoforms on the level of single native RNA molecules. nano-ID combines metabolic RNA labeling with native RNA nanopore sequencing to enable RNA isoform identification, estimation of its stability, and a measurement of its poly(A)-tail length from a single sample. nano-ID is able to visualize changes in RNA isoform synthesis and stability and reveals a hidden layer of gene regulation. nano-ID thus allows to study transcriptional regulation in unprecedented detail and can prevent misleading conclusions that would be obtained when only combined RNAs from an entire gene locus are considered, as is done by RNA-seq, 4sU-seq or TT-seq.

nano-ID has many advantages over other sequencing-based transcriptomic strategies as it allows to sequence the original native RNA molecule. In particular, there is no need for fragmentation of RNA prior to sequencing and hence no ambiguity in assigning reads to RNA isoforms. nano-ID also does not require template amplification and thus omits copying errors and sequence-dependent biases. It comes without a lengthy library protocol and eliminates sequencing by synthesis and therefore prevents loss of information on epigenetic modifications and artificially introduced RNA base analogues. It is PCR-free and shows neither sequence bias nor read duplication events. Taken together, it overcomes drawbacks and limitations of state-of-the-art approaches and increases the gathered information vastly.

Generally, nanopore sequencing has still limitations in throughput and accuracy. These drawbacks, however, are outweighed by the information obtained on the sequencing substrates. The longer the sequenced molecules are, the less problematic is the lack in accuracy in identifying their origin or classifying it into newly-synthesized or pre-existing. On top of that, there are strategies to improve splice site calling with already existing high accuracy ‘short-read’ sequencing data to reduce sequencing errors or to assess the likelihood of real nucleotide variants. We can however show that our algorithmic strategies are already sufficient to address metabolic rate estimation in a reliable manner. Technical improvements in nanopore sequencing or their computational processing will strongly improve the accuracy of individual read sequences and thus detectability of ^5E^U. The task at hand will be the development of a novel base-calling algorithm for direct RNA nanopore sequencing with extended base alphabet (A, C, G, U & ^5E^U). Furthermore, increased throughput will foster statistical precision of metabolic rate estimation and will also allow to elucidate low abundant or transient processes.

Nanopore-based transcriptomic studies will allow us to monitor the formation of transcripts, post-transcriptional processing, export and translation at the level of single RNA isoforms. nano-ID is in principle also transferable to single cell methodologies, to catch heterogeneity of the RNA population in any state of the cell. This however requires sequencing library preparation with lower input amounts. The use of ^5E^U is widely established for *in vivo* applications in the field such as fluorescence microscopy. We thus envision that nano-ID is in principle applicable to many types of organisms, cells and conditions.

## Methods

### Labeling and direct RNA nanopore sequencing of synthetic RNAs

Labeled synthetic RNAs and synthetic control RNAs are derived from selected RNAs of the ERCC RNA Spike-in Mix (Ambion) as described in ^26^. Characteristics of selected RNAs of the ERCC RNA Spike-in Mix are listed in (**Supplementary Table 3**). Briefly, selected spike-in sequences were cloned into a pUC19 cloning vector and verified by Sanger sequencing. For IVT template generation, the plasmid (3 µg) was linearized using EcoRV-HF (blunt end cut) digestion mix containing CutSmart buffer and EcoRV-HF enzyme. The digestion mix was incubated at 37 °C for 1 h and the reaction was terminated adding 1/20 volume of 0.5 M EDTA. Subsequently, DNA was precipitated in 1/10 volume of 3 M sodium acetate pH 5.2, and 2 volumes of 100 % ethanol at - 20 °C for 15 min. DNA was collected by centrifugation at 4 °C and 16,000 x g for 15 min. The pellet was washed twice using 75 % ethanol. DNA was air-dried and resuspended in 5 µL of H_2_O at a concentration of 0.1-1.0 µg/µL (quantified by NanoDrop). Synthetic RNAs were *in vitro* transcribed using the MEGAscript T7 kit (Ambion). *In vitro* transcription (IVT) of synthetic control RNAs was performed following the manufacturer’s instruction. For IVT of labeled synthetic RNAs, 100 % of UTP (resp. GTP) was substituted with either 5-ethynyl-UTP (^5E^U, Jena Bioscience), 5-bromo-UTP (^5Br^U, Sigma), 5-iodo-UTP (^5I^U, TriLink BioTechnologies LLC), 4-thio-UTP (^4S^U, Jena Bioscience) or 6-thio-GTP (^6S^G, Sigma). Note that, for performing a successful IVT with 4-thio-UTP and 6-thio-GTP, only a reduction to 80% substitution gave successful yield. IVT reactions were incubated at 37 °C. After 4 h, reaction volume was filled up with H_2_O to 40 µL, then 2 µL of TURBO DNase was added and incubated at 37 °C for additional 15 min. Synthetic RNAs were purified with RNAClean XP beads (Beckman Coulter) following the manufacturer’s instructions. The final synthetic RNA pool contained equal mass of all respective synthetic RNAs in a given library (**Supplementary Table 1**). RNA was quantified using Qubit (Invitrogen). RNA quality was assessed with the TapeStation System (Agilent) Synthetic RNA pools were poly(A)-tailed using the *E. coli* Poly(A) Polymerase (NEB). The reaction was incubated for 5 min and stopped with 0.1 M EDTA. Spike-ins were then purified with phenol:chloroform:isoamyl alcohol and precipitated. Poly(A)-tailed synthetic RNA pools were subsequently subjected to direct RNA nanopore sequencing library preparation (SQK-RNA001, Oxford Nanopore Technologies) following manufacturer’s protocol. All libraries were sequenced on a MinION Mk1B (MIN-101B) for 20 h, unless reads sequenced per second stagnated dramatically.

### Culturing of human K562 cells

Human K562 erythroleukemia cells were obtained from DSMZ (Cat. # ACC-10). K562 cells were cultured antibiotic-free in accordance with the DSMZ Cell Culture standards in RPMI 1640 medium (Thermo Fisher Scientific) containing 10 % heat inactivated fetal bovine serum (FBS) (Thermo Fisher Scientific), and 1x GlutaMAX supplement (Thermo Fisher Scientific) at 37 °C in a humidified 5 % CO_2_ incubator. Cells used in this study display the phenotypic properties, including morphology and proliferation rate, that have been described in literature. Cells were verified to be free of mycoplasma contamination using Plasmo Test Mycoplasma Detection Kit (InvivoGen). Biological replicates were cultured independently.

### ^5E^U labeling and direct RNA nanopore sequencing of human K562 cells

K562 cells were kept at low passage numbers (<6) and at optimal densities (3×10^5 - 8×10^5) during all experimental setups. Per biological replicate, K562 cells were diluted 24 h before the experiment was performed (**Supplementary Table 1**). Per ^5E^U 60 min sample (4 replicates), cells were incubated at 37 °C, 5 % CO_2_ for 1 h after a final concentration of 500 µM 5-Ethynyluridine (^5E^U, Jena Bioscience) was added. Per ^5E^U 24 h sample (3 replicates), cells were incubated at 37 °C, 5% CO_2_ for 24 h. ^5E^U was added 3 times during the 24h incubation, i.e. every 8 hours (0h, 8h, 16h) at a final concentration of 500 µM. Control samples were not labeled (3 replicates). Per ^5E^U 60 min HS (heat shock) sample (3 replicates), cells were incubated at 42 °C for 5 min (until cell suspension reached 42 °C), and then ^5E^U was added at a final concentration of 500 µM. Further, heat shock treatments were performed in a water bath (LAUDA, Aqualine AL12) at 42 °C. for 1 h. Temperature was monitored by thermometer. To avoid transcriptional changes by freshly added growth medium, fresh growth medium was added ∼24 h prior to heat shock treatments ^43^. Exactly after the labeling duration, cells were centrifuged at 37 °C and 1,500 x g for 2 min. Total RNA was extracted from K562 cells using QIAzol (Quiagen) according to manufacturer’s instructions. Poly(A) RNA was purified from 1 mg of total RNA using the µMACS mRNA Isolation Kit (Milteny Biotec) following the manufacturer’s protocol. The quality of poly(A) RNA selection was assessed using the TapeStation System (Agilent). Poly(A) selected RNAs were subsequently subjected to direct RNA nanopore sequencing library preparation (SQK-RNA001, Oxford Nanopore Technologies) following manufacturer’s protocol with 1000 ng input. All libraries were sequenced on a MinION Mk1B (MIN-101B) for 48 h, unless reads sequenced per second stagnated dramatically.

### RNA-seq

Two biological replicates of K562 cells were diluted 24 h before the experiment was performed. Per replicate, 3.6 x 10^7^ cells in growth medium were labeled at a final concentration of 500 µM 4-thio-uracil (4sU, Sigma-Aldrich), and incubated at 37 °C, 5 % CO_2_ for 5 min. Exactly after 5 min of labeling, cells were harvested at 37 °C and 1,500 x g for 2 min. Total RNA was extracted from K562 cells using QIAzol according to manufacturer’s instructions except for the addition of 150 ng RNA spike-in mix ^26^ together with QIAzol. To isolate polyA RNA from 75 µg of total RNA, two subsequent rounds of purification by Dynabeads Oligo (dT)_25_ (invitrogen) were performed. Purification based on manufacturer’s instructions was performed twice, using 1 mg of Dynabeads Oligo (dT)_25_ beads for the first round and 0.5 mg for the second round of purification. The quality of polyadenylated RNA selection was assessed using RNA ScreenTape on a TapeStation (Agilent). Sequencing libraries were prepared using the NuGEN Ovation Universal RNA-seq kit according to manufacturer’s instructions. Fragments were amplified by 10 cycles of PCR, and sequenced on an Illumina NextSeq 550 in paired-end mode with 75 bp read length.

### Direct RNA nanopore sequencing data preprocessing of synthetic RNAs

Direct RNA nanopore sequencing reads were obtained for each of the samples (**Supplementary Table 1**). FAST5 files were base-called using Albacore 2.3.1 (Oxford Nanopore Technologies) with the following parameters: read_fast5_basecaller.py -f FLO-MIN106 -k SQK-RNA001. Direct RNA nanopore sequencing reads were mapped with GraphMap 0.5.2 ^44^ to the synthetic RNA reference sequence with the following parameters: graphmap align --evalue 1e-10. Further data processing was carried out using the R/Bioconductor environment.

### Direct RNA nanopore sequencing data preprocessing of human K562 cells

Direct RNA nanopore sequencing reads were obtained for each of the samples (**Supplementary Table 1**). FAST5 files were base-called using Albacore 2.3.1 (Oxford Nanopore Technologies) with the following parameters: read_fast5_basecaller.py -f FLO-MIN106 -k SQK-RNA001. Direct RNA nanopore sequencing reads were mapped with Minimap2 2.10 ^45^ to the hg20/hg38 (GRCh38) genome assembly (Human Genome Reference Consortium) with the following parameters: minimap2 -ax splice -k14 --secondary=no. Samtools ^46^ was used to quality filter SAM files, whereby alignments with MAPQ smaller than 20 (-q 20) were skipped. Further data processing was carried out using the R/Bioconductor environment and custom python scripts.

### Probability of ^5E^U-labeled RNA isoform identification based on synthetic RNAs

The following parameters were collected on the direct RNA nanopore sequencing data of synthetic RNAs and used to calculate the probability of identification of a ^5E^U-labeled RNA isoform as labeled. Detectability *d* - the number of ^5E^U caused mismatches in the ^5E^U-labeled sample. Background *b* - the number of U caused mismatches in the unlabeled control sample. Given these parameters, the probability of identification *p* can be calculated as the probability of a U-based mismatch being caused by a ^5E^U in the transcript as

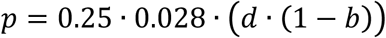

with *0.25* - the empirical probability of U content, and labeling efficiency *0.028* - the empirical probability of a U being replaced by a ^5E^U in the labeling process ^32^. This then allows to calculate the probability of labeled RNA identification *p*^*RNA*^ as

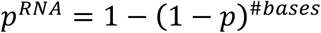

, the probability, that an RNA contains at least 1 detectable ^5E^U.

### Definition of transcription units based on the UCSC RefSeq genome assembly GRCh38 (RefSeq-TUs)

For each annotated gene, transcription units were defined as the union of all existing inherent transcript isoforms (UCSC RefSeq GRCh38).

### Definition of isoform-independent exonic and intronic regions (constitutive exons and introns)

Isoform-independent exonic and intronic regions were determined using a model for constitutive exons ^47^ and constitutive introns respectively based on UCSC RefSeq annotation (GRCh38).

### Isoform determination for human K562 cells

The FLAIR (Full-Length Alternative Isoform analysis of RNA) algorithm ^22^ was used for the correction and isoform definition of raw human K562 direct RNA nanopore reads. Corrected and collapsed isoforms were obtained by adding short-read data (RNA-seq) to help increase splice site accuracy of the nanopore read splice junctions (https://github.com/BrooksLabUCSC/FLAIR).

### Parameter collection for neural network training and classification

For each read in each human K562 sample (^5E^U 60 min, Control, ^5E^U 24 h & ^5E^U 60 min HS) we obtained ∼1500 parameters from three different layers: Raw signal (ionic current), base-call event probabilities and alignment derived mismatch properties. As raw signal, 1193 parameters were gathered consisting of the raw ionic current measurements gathered for each possible 5-mer of nucleotides as well as the raw ionic current measurements gathered for each possible 3-mer centered in a 5-mer. The latter parameters were collected for U-containing and non-U-containing instances. In addition to that, raw ionic current measurements were gathered for 5-mers with all possible nucleotides in their center position also for U-containing and non-U-containing instances, as well as 5-mers exclusively leading or lagging U content. All collected raw signal parameters were z-score normalized on all non-U-containing instances given the mean values of the pore model on which the original base-calling algorithm is based provided by Oxford Nanopore Technologies. As base-call event probabilities, 120 parameters were gathered including ‘model state’, ‘move’, ‘weights’, ‘p model state’, the probability that ‘model state’ gave rise to the observation of the event, the most probable ‘model state’, the probability that ‘p model state’ gave rise to the observation of the event and the probabilities that events may be associated with the certain base from the event probabilities table provided by the base-calling algorithm. As alignment derived mismatch properties, 135 parameters were gathered including length of the reads, nucleotide occurrences, number of nucleotide transitions, number of inserts and deletions on a single nucleotide basis as well as on a 5-mer basis for U-containing and non-U-containing instances.

### Neural network training, validation and classification of human RNA isoforms into ^5E^U-labeled and unlabeled

Neural network was trained on the ^5E^U 24 h versus Control samples under the assumption that ^5E^U 24 h sample solely contains labeled reads and the fact that the Control sample solely contains unlabeled reads. The trained neural network consists of a batch normalization layer and 3 dense layers with decreasing output shape (**Supplementary Figure 3a**). 2 dropout layers (with 25% dropout) in between regularize the attempted classification. Training was conducted on 404.201 reads, validation was performed on 173.240 reads in 40 epochs with the R interface to Keras on a TensorFlow backend ^48^, as

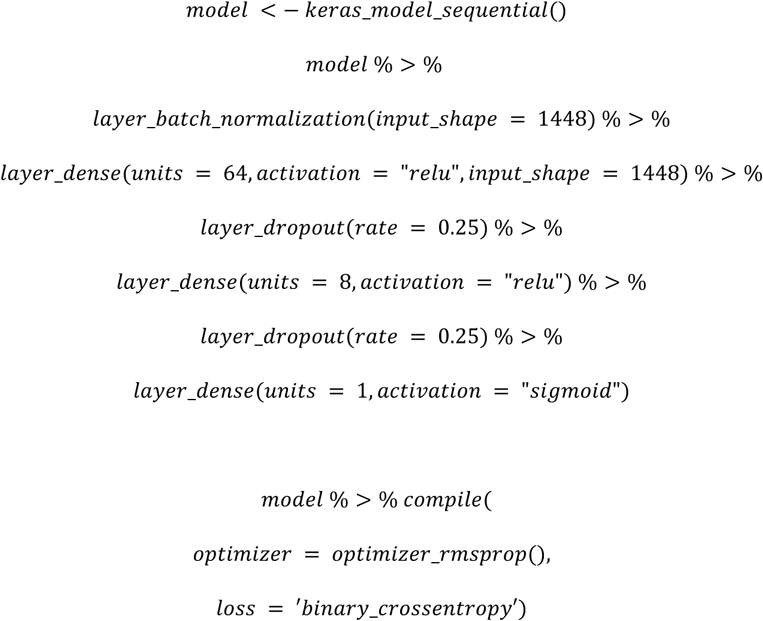

The neural network was 5-fold cross-validated with an accuracy of 0.87 and a false discovery rate (FDR) of 0.025 and used to classify reads of the ^5E^U 60 min and ^5E^U 60 min HS samples into ^5E^U-labeled and unlabeled. A ROC analysis (1 – specificity vs sensitivity) for all reads of the test set showed an area under the curve (AUC) of 0.94. For reads with an alignment length larger than 500 nt and 1000 nt the AUC improved to 0.96. Note that, limiting the neural network classification to reads produced in the first few hours of sequencing, i.e. reads with a generally higher accuracy, improves the AUC to 0.98.

### Random forest training, validation and classification of human RNA isoforms into ^5E^U-labeled and unlabeled

For validation purposes, a random forest ^49^ was trained on the ^5E^U 24 h versus Control samples on the same data as the neural network above. The random forest was 5-fold cross-validated with an accuracy of 0.85 and a false discovery rate (FDR) of 0.32 and used to classify reads of the ^5E^U 60 min sample into ^5E^U-labeled and unlabeled.

### Poly(A)-tail length determination

Poly(A)-tail length is estimated by identifying the dwell time of the poly(A)-tail in the nanopore. For each direct RNA nanopore sequencing read, the raw signal readout of the nanopore in pico-Ampere [pA] was extracted from the FAST5 file. Every data point above the 99.99% quantile or below the 0.001% quantile was set to the respective cut-off value for reasons of robustness (**Supplementary Figure 5c, upper panel**). Subsequently kmeans clustering was used to define two trend lines at 1/3 and 2/3 the distance between the two cluster centers. The two trend lines were then used to squish the raw data by taking the parallel minimum or maximum (**Supplementary Figure 5c, lower panel**). A loss score of a piecewise linear function of two consecutive segments of the trend lines is then used to identify segments along the squished data points (**Supplementary Figure 5c, middle panel**). The length of the third identified segment *r*_*j*_ is used to calculate the length of the poly(A)-tail *l*_*j*_ of read *j* in sample *i* as

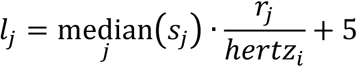

with the sequencing read speed *s*_*j*_ of read *j* in [nt/s] and the frequency *hertz*_*i*_ in [Hz] used in measuring sample *i* and 5 additional adenines that are concealed in the flanking 5-mers.

### Intron retention ratio

For each RefSeq-TU (UCSC RefSeq GRCh38) the intron retention ratio for the ^5E^U 60 min and ^5E^U 60 min HS samples were calculated using the above defined model of constitutive exons and introns by calculating the ratio of length normalized coverages of the maximum value for all respective introns and the average of all respective exons. This yielded 358 gene loci with at least 5% intron retention in either of the samples.

### RNA stability (degradation rate *λ*_*ij*_, half-life *hl*_*ij*_) and synthesis rate *μ*_*ij*_ estimation of human RNA isoforms

Each neural network classified direct RNA nanopore sequencing read of the ^5E^U 60 min and ^5E^U 60 min HS samples was assigned to a FLAIR defined human isoform (or RefSeq-TU) either as ^5E^U-labeled *L*_*ij*_ and unlabeled *T*_*ij*_ *– L*_*ij*_. The resulting counts were subsequently converted into synthesis rates *μ*_*ij*_ and degradation rates *λ*_*ij*_ for isoform *i* in sample *j* assuming first-order kinetics as in ^24^ using the following equations:

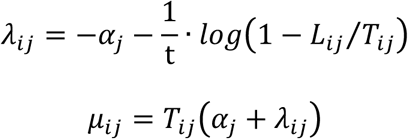

where *t* is the labeling duration in minutes and *α* is the growth rate (dilution rate, i.e. the reduction of concentration due to the increase of cell volume during growth) defined as

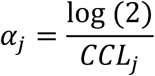

with cell cycle length *CCL*_*j*_ [min]. The half-life *hl*_*ij*_ for isoform *i* in sample *j* can thus be calculated as

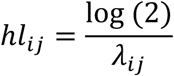

in minutes [min].

### RNA-seq data preprocessing and antisense bias correction

Paired-end 75 base reads with additional 6 base reads of barcodes were obtained for each of the samples (**Supplementary Table 1**). Reads were demultiplexed and mapped with STAR 2.3.0 ^50^ to the hg20/hg38 (GRCh38) genome assembly (Human Genome Reference Consortium). Samtools ^46^ was used to quality filter SAM files, whereby alignments with MAPQ smaller than 7 (-q 7) were skipped and only proper pairs (-f2) were selected. Further data processing was carried out using the R/Bioconductor environment. We used a spike-in (RNAs) normalization strategy essentially as described ^26^ to allow observation of antisense bias ratio *c*_*j*_ (ratio of spurious reads originating from the opposite strand introduced by the reverse transcription reaction). Antisense bias ratios were calculated for each sample *j* according to

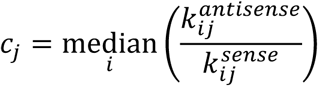

for all available spike-ins *i*. Read counts (*k*_*ij*_) for spike-ins were calculated using HTSeq ^51^. The number of transcribed bases (*tb*_*j*_) for all samples was calculated as the sum of the coverage of evident (sequenced) fragment parts (read pairs only) for all fragments in addition to the sum of the coverage of non-evident fragment parts for fragments with an inner mate interval not entirely overlapping a Refseq annotated intron (UCSC RefSeq GRCh38). The number of transcribed bases (*tb*_*j*_) or read counts (*k*_*j*_) for all features (RefSeq-TUs) were corrected for antisense bias *c*_*j*_ as follows using the parameter calculated as described above. The real number of read counts or coverage *s*_*ij*_ for transcribed unit *i* in sample *j* was calculated as

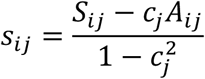

where *S*_*ij*_ and *A*_*ij*_ are the observed numbers of read counts or coverage on the sense and antisense strand. RPKs were calculated upon antisense bias corrected read counts (*k*_*j*_) falling into the region of a RefSeq-TU divided by its length in kilobases. Coverages were calculated upon antisense bias corrected number of transcribed bases (*tb*_*j*_) falling into the region of a RefSeq-TU divided by its length in bases.

## Supporting information

nano-ID gene level estimates

nano-ID isoform level estimates

## Acknowledgments

We would like to thank Johannes Söding, Christian Roth and Dmitry Tegunov (Max Planck Institute for Biophysical Chemistry) for advice on machine learning aspects. We would also like to thank Nikolaos Papadopoulos and Noah Wulff Mottelson (Max Planck Institute for Biophysical Chemistry) for help with python. In addition, we would like to thank Michael Lidschreiber (Max Planck Institute for Biophysical Chemistry) and Brian Lee (UCSC Genomics Institute) for help with the UCSC Genome Browser Track Hub. Moreover, we would like to thank Anna Sawicka and Kristina Žumer (Max Planck Institute for Biophysical Chemistry) for sharing the pUC19 spike-in plasmids. PC was supported by the Deutsche Forschungsgemeinschaft (SFB860, SPP1935), the European Research Council Advanced Investigator Grant TRANSREGULON (grant agreement No 693023), and the Volkswagen Foundation.

## Competing interests

The authors declare that no competing interests exist.

## Authors’ contributions

KM, BS and SG carried out experiments. BS designed and carried out all bioinformatics analysis. BS conceptualized, designed and supervised research. BS and PC prepared the manuscript, with input from all authors.

## Notes

#### Summary of Updates

Additional supplementary *.txt files have been provided giving the isoform and gene level estimates of nano-ID, such as synthesis rates, half-lives and poly(A)-tail lengths.

