## Supplementary Information for "Native molecule sequencing by nano-ID reveals synthesis and stability of RNA isoforms"

Supplementary note 1 ........................................ p. 1
Supplementary figures 1-7 ........................................ p. 2-10
Supplementary tables 1-5 ........................................ p. 11-15
References ........................................ p. 16

### Supplementary note

#### Supplementary Note 1. General and method-specific considerations.

$^{5E}$U uptake and incorporation efficiency might vary between cell-types and organisms, and thus a careful assessment of $^{5E}$U-labeling conditions and exposure time is highly recommended as it can influence the ability of metabolic rate estimation. For studying intracellular RNA isoform kinetics, it is statistically speaking of great interest for the whole approach to keep cell exposure times to a feasible minimum. It is worth noting that $^{5E}$U is incorporated *in vivo* with an efficiency (~2-3%) that is unlikely disruptive to the physiology of the cell at short exposure times.

### Supplementary figures

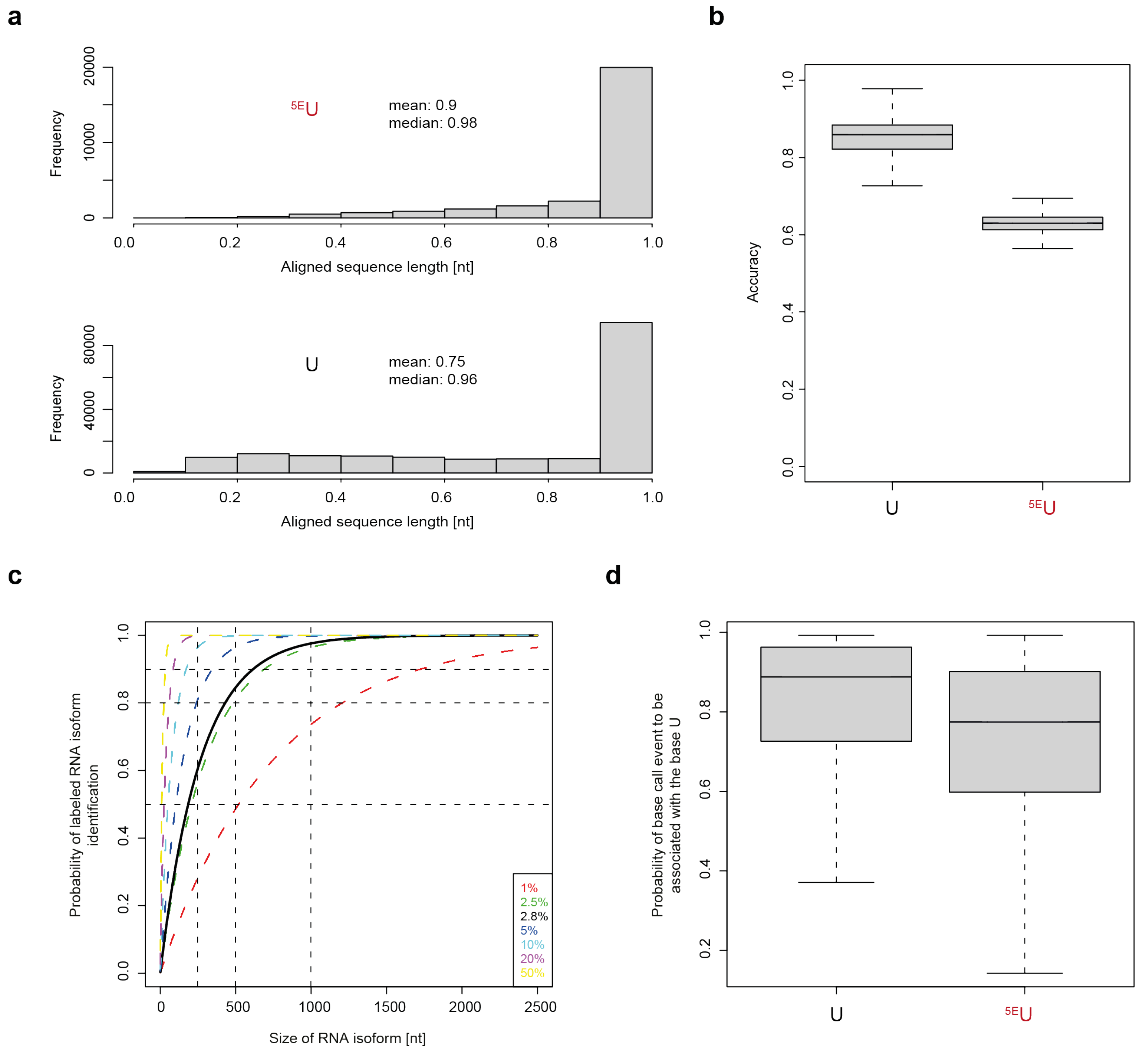


**Supplementary Figure 1. Full-length and accuracy assessment of synthetic RNAs and probability of $^{5E}$U-labeled RNA isoform identification.** (a) Histograms show the relative length of aligned reads of synthetic RNAs containing $^{5E}$U instead of U ($^{5E}$U, 23,805 molecules, mean: 0.9, median: 0.98) and synthetic control RNAs (U, 175,389 molecules, mean: 0.75, median: 0.96) relative to the reference sequence. (b) Boxplot shows the read based accuracy (edit distance) of aligned synthetic RNAs containing $^{5E}$U instead of U ($^{5E}$U, 23,805 molecules) and synthetic control RNAs (U, 175,389 molecules) based upon the reference sequence. (c) Plot shows the probability of $^{5E}$U-labeled RNA isoform identification dependent on alignment length [nt] per RNA isoform and labeling efficiency (**Methods**). (d) Boxplot shows the probability of base-called U or $^{5E}$U instances in synthetic RNAs containing $^{5E}$U instead of U ($^{5E}$U, 23,805 molecules) and synthetic control RNAs (U, 93,060 molecules) to be associated with the base U calculated by standard base-calling algorithm (right).

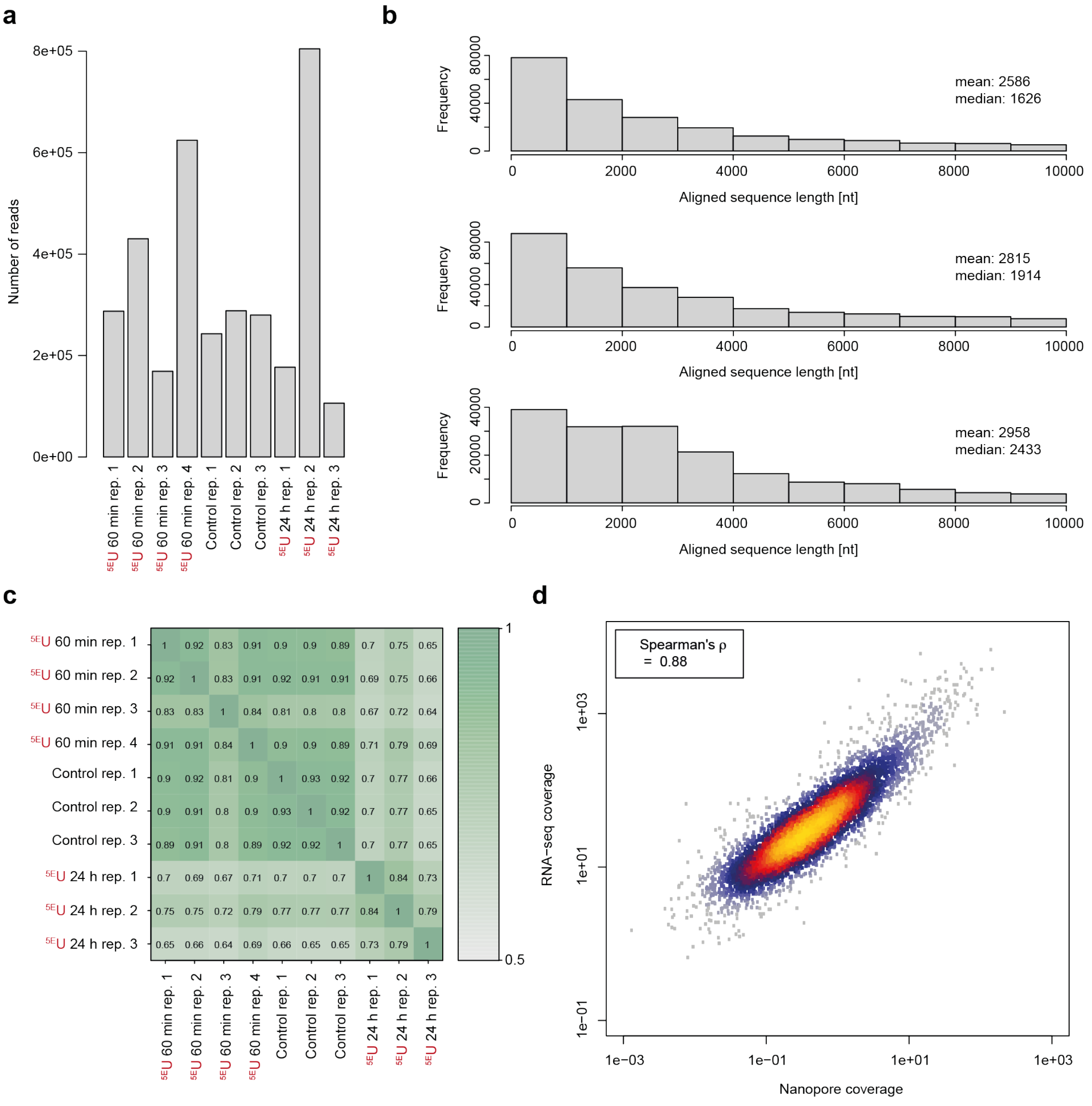


**Supplementary Figure 2. Quality metrics and reproducibility assessment of direct RNA 'long-read' nanopore sequencing of RNA isoforms in human K562 cells.** (a) Barplot shows the obtained number of reads for all samples (4 biological replicates $^{5E}$U 60 min, 3 biological replicates Control & 3 replicates $^{5E}$U 24 h). (b) Histograms show the length distribution of aligned reads of human RNA isoforms. From top to bottom: 3 replicates $^{5E}$U 60 min (218,755 molecules, mean: 2,586 nt, median: 1,626 nt), 3 replicates Control (280,435 molecules, mean: 2,815 nt, median: 1,914 nt) & 3 replicates $^{5E}$U 24 h (167,317 molecules, mean: 2,958 nt, median: 2,433 nt). (c) Heatmap showing pairwise Spearman correlation coefficients of RefSeq GRCh38 annotation based read counts of all replicates in all samples. Color code indicates Spearman correlation coefficients. Coefficients shown are rounded to the second decimal place. (d) Scatter plot with color-coded density of read coverage of 4 biological replicates $^{5E}$U 60 min and 3 biological replicates Control (Nanopore) versus read coverage of two Illumina sequencing based replicates of poly(A)-selected RNAs (RNA-seq, **Methods**). Shown are 6,458 RefSeq GRCh38 annotated genes with at least 2 reads in all direct RNA 'long-read' nanopore sequencing samples. Correlation is calculated as Spearman's rank correlation coefficient (0.88) rounded to the second decimal.

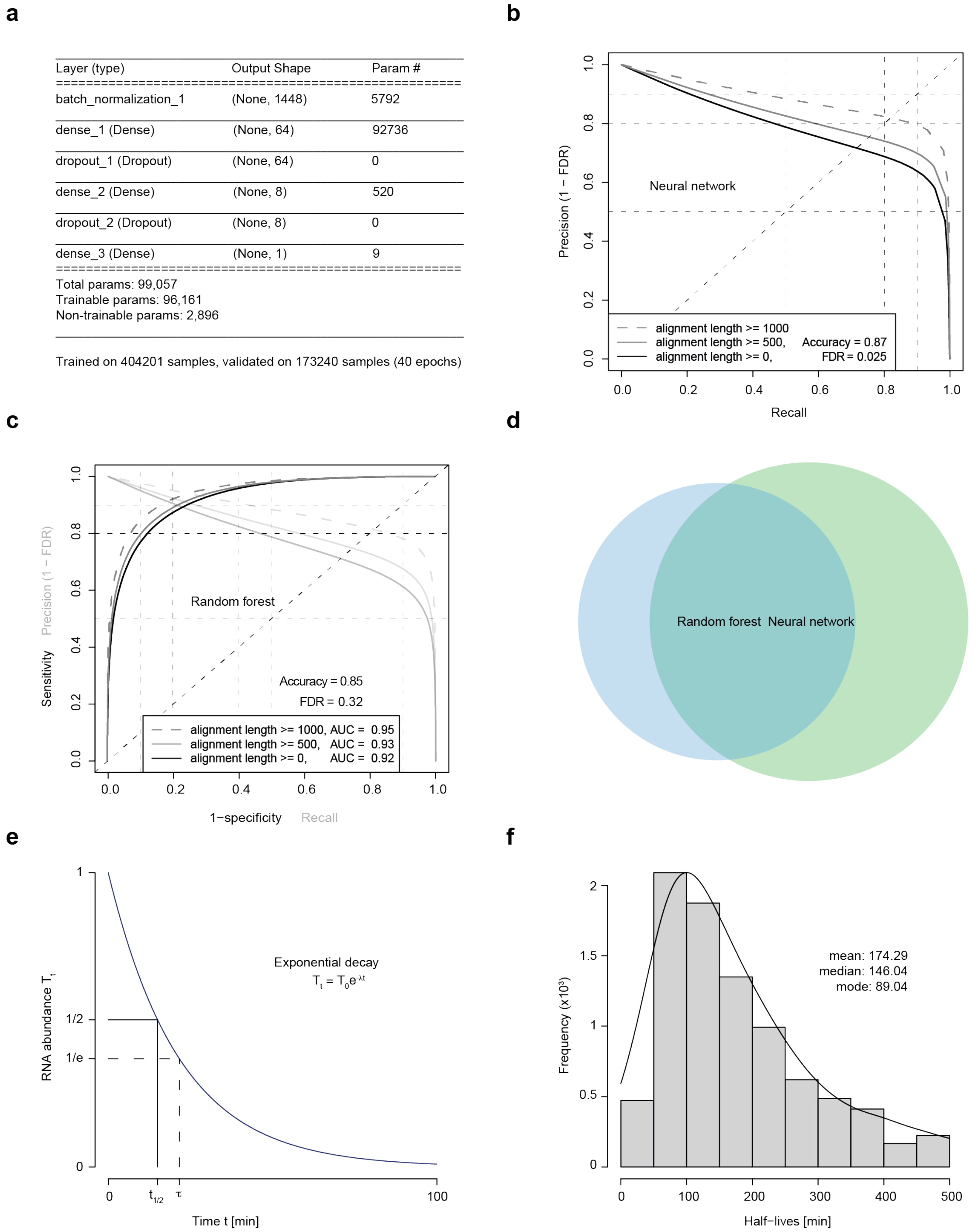


**Supplementary Figure 3. Design and quality metrics of neural network based classification of human RNA isoforms into $^{5E}$U-labeled and unlabeled.** (a) Layer design of neural network. It consists of a batch normalization layer and three dense layers with decreasing output shape. Two dropout layers in between regularize the attempted classification. Training was conducted on 404,201 reads, validation was performed on 173,240 reads in 40 epochs. (b) Precision and recall analysis of 5-fold cross-validated neural network training. Plot shows precision versus recall curves for all reads of the test set (black, alignment length >=0), for reads with an alignment length larger than 500 nt (grey, alignment length >=500) and for reads with an alignment length larger than 1,000 nt (dashed grey, alignment length >=1000). (c) ROC analysis of 5-fold cross-validated random forest training (**Methods**) as in (**Fig. 4b**) with additional precision and recall analysis as in (b) depicted with lower opacity. (d) Euler diagram comparing the number of reads classified as $^{5E}$U-labeled by the neural network (light green) and by the random forest approach (light blue). The overlap is 53% and 70%, respectively. (e) Schematic illustration of exponential decay model (Methods) used to calculate half-lives of RNAs $\lambda$ (mean lifetime $\tau$). (f) Histogram of RNA half-life estimates of 8,687 RefSeq-TUs (mean: 174 min, median: 146 min).

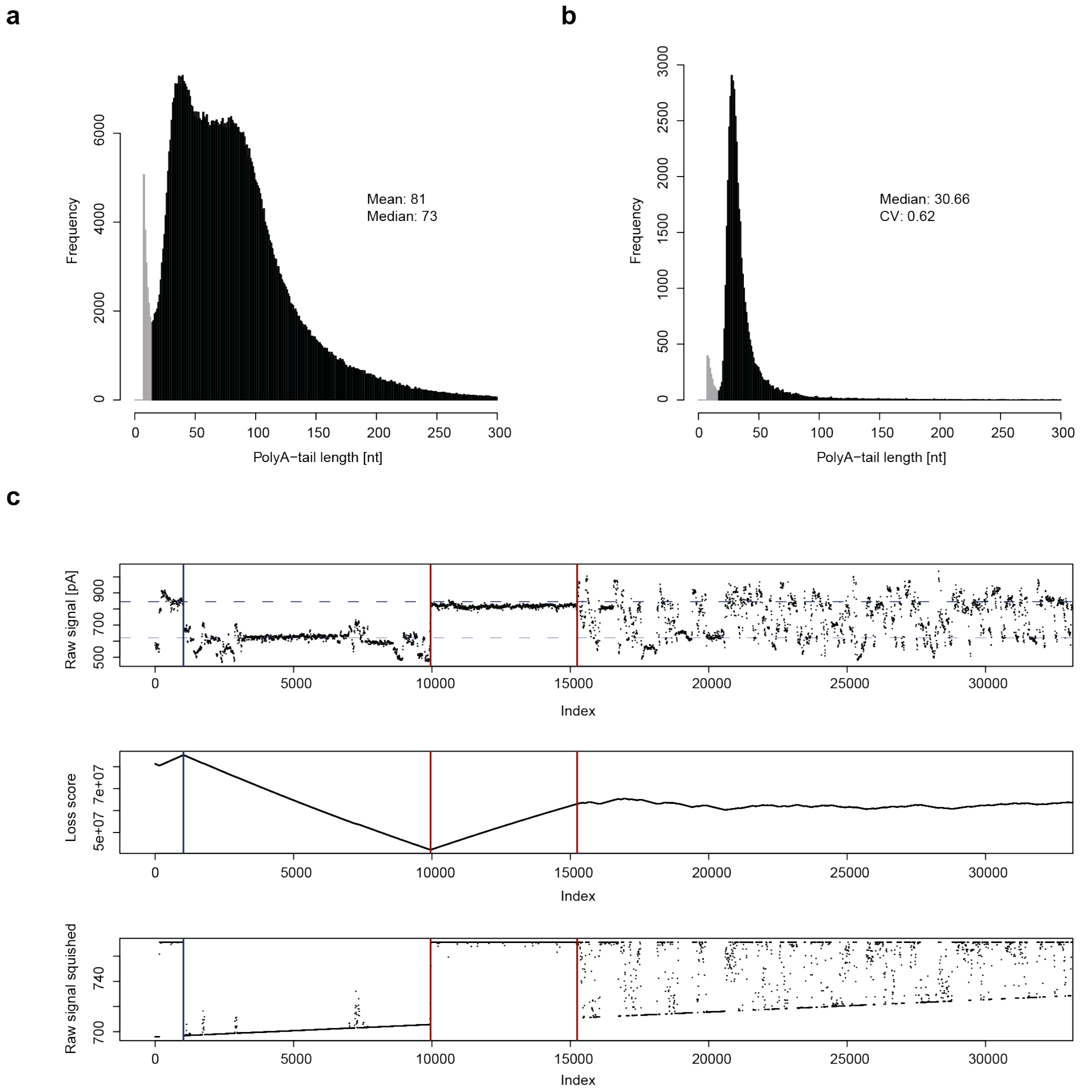


**Supplementary Figure 4. Detailed view on poly(A)-tail length determination of human RNA isoforms.** (a) Histogram of poly(A)-tail length estimates of 714,536 RNA isoforms using 300 bins (mean: 81 nt, median: 73 nt). (b) Histogram of poly(A)-tail length estimates of the RNA calibration strand (RCS) using 300 bins (median: 30.6, coefficient of variation: 0.62). (c) Schematic view on poly(A)-tail length determination. Upper panel: raw signal readout of the nanopore in pico-Ampere [pA] of exemplary read. Poly(A)-tail length is estimated by identifying the dwell time of the poly(A)-tail in the nanopore (depicted between two vertical red lines) followed by the signature of the sequencing adapter (depicted before the first vertical red line). Middle panel: loss score of piecewise linear function used to identify segments in the raw signal readout. Lower panel: squished raw signal (given an upper and lower bound) which introduces robustness into poly(A)-tail length identification.

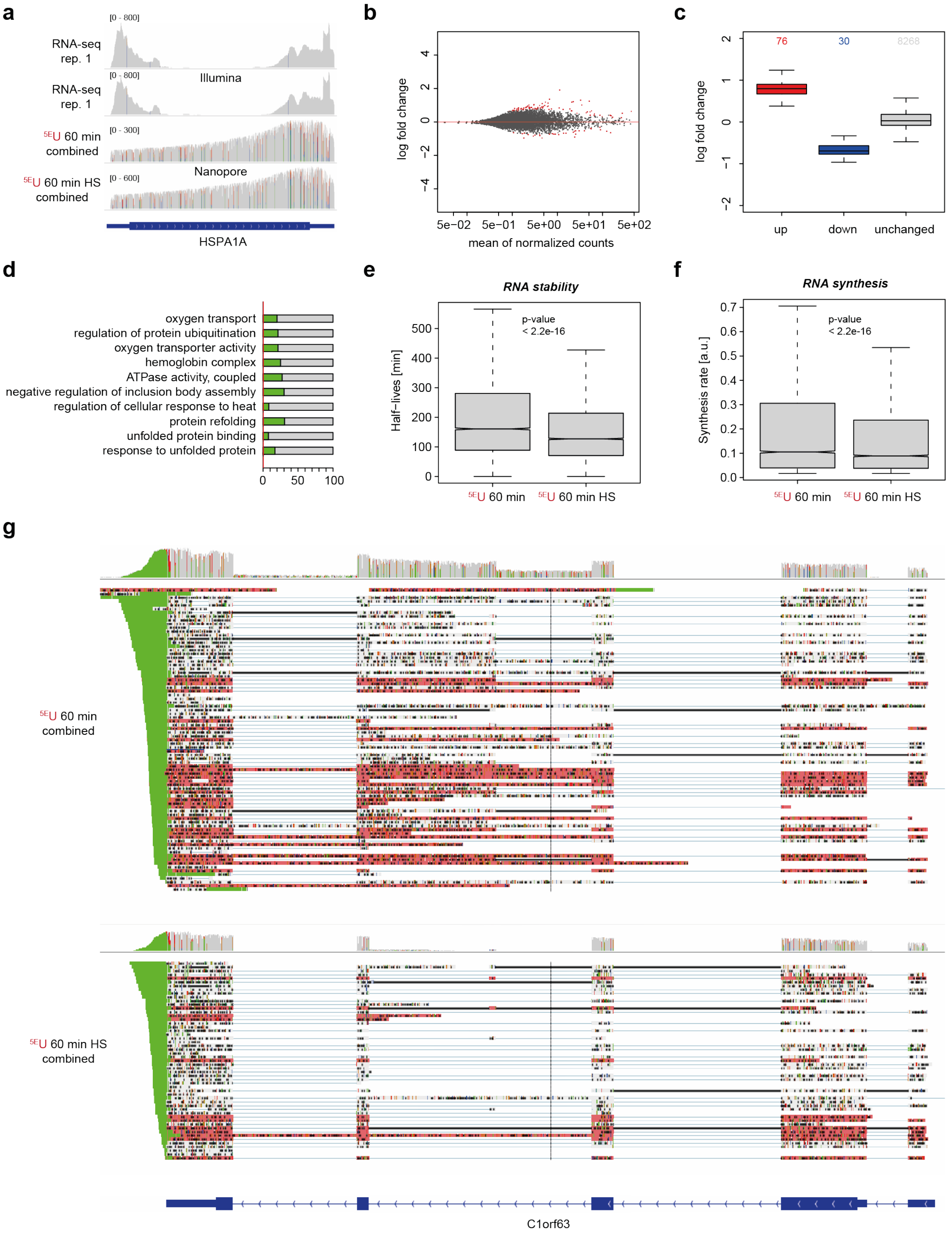


**Supplementary Figure 5. nano-ID captures the response to heat shock in human K562 cells and reveals the biogenesis of RNA isoforms.** (a) Genome browser view of the human HSPA1A gene locus (chr6:31,813,514-31,819,942). ‘Long-read’ nanopore sequencing (lower panels) resolves mappability problems of RNA-seq (lower panels). (b) Log fold change upon heat shock treatment versus the normalized mean read count of $^{5E}$U-labeled reads. Heat shock samples ($^{5E}$U 60 min HS) were compared to respective control ($^{5E}$U 60 min). Significantly up- or downregulated RefSeq-annotated genes (p-value < 0.1) are marked in red. (c) Boxplot shows upregulated (red), downregulated (blue) and unchanged RefSeq annotated genes (grey) in human K562 cells upon 60 min of heat shock (42 °C). A minimum fold change of 1.25 and a maximum p-value of 0.1 was set for calling a significant expression change. (d) Gene Ontology (GO) analysis [1] of significantly overrepresented categories linked to upregulated RefSeq-annotated genes upon heat shock for human K562 cells. Red line, proportion of upregulated RefSeq annotated genes in the whole population. The number of upregulated RefSeq annotated genes in the resp. GO category is given relative to the GO category size (green bar). (e) Boxplot shows half-lives of RNAs comparing heat shock ($^{5E}$U 60 min HS) against control ($^{5E}$U 60 min). (f) Boxplot shows synthesis rate of RNAs comparing heat shock ($^{5E}$U 60 min HS) against control ($^{5E}$U 60 min). (g) Genome browser view of classified direct RNA nanopore sequencing reads of the human C1orf63 gene locus on chromosome 1 (~5.6 kbp, chr1: 25,241,897-25,247,552) comparing heat shock ($^{5E}$U 60 min HS) against control ($^{5E}$U 60 min) visualized with the Integrative Genomics Viewer (IGV, version 2.4.10; human hg38) [2]. Unlabeled reads are shown in grey, $^{5E}$U-labeled reads are shown in red, and poly(A)-tails are highlighted in green.

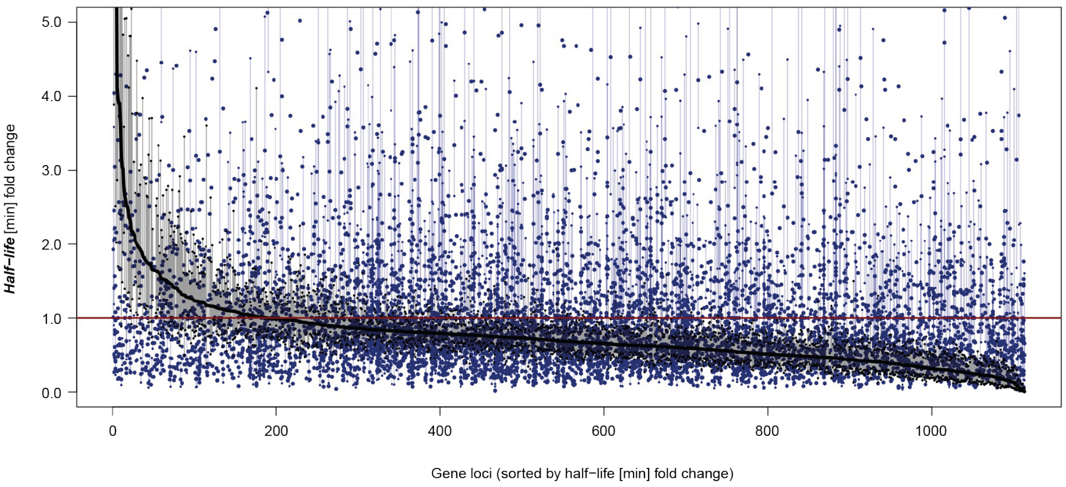


**Supplementary Figure 6. nano-ID allows full dynamic characterization of human RNA isoforms.** Half-life fold change (y-axis) depicted across 1,113 gene loci (x-axis). Half-life estimates for entire gene loci (combined) are depicted as a black line (sorted in decreasing order). Blue dots represent individual isoform half-life estimates at respective gene loci (7,627 isoforms in total). All estimates are supported by at least 1 biological replicate in each condition. Perpendicular blue and black lines represent standard deviations of individual estimates. For individual RNA isoform half-life estimates, standard deviations are only shown if not overlapping with the standard deviation of the respective combined half-life estimates (black).

### Supplementary tables

**Supplementary Table 1. Information on experimental conditions used in this study.** Abbreviations used: $^{5E}$U for 60 minutes ($^{5E}$U 60 min), 24 h ($^{5E}$U 24 h, heat shock (HS) and direct RNA 'long-read' nanopore sequencing (Nanopore).

| No. | Assay | Cell type | Condition name | Replicate no. | Treatment |
|---|---|---|---|---|---|
| **1** | Nanopore | - | Synthetic control | 1 | IVT |
| **2** | Nanopore | - | $^{5Br}$U, $^{4S}$U, $^{6S}$G | 1 | $^{5Br}$U, $^{4S}$U, $^{6S}$G, IVT |
| **3** | Nanopore | - | $^{5E}$U, $^{5I}$U | 1 | $^{5E}$U, $^{5I}$U, IVT |
| **4** | Nanopore | K562 | $^{5E}$U 60 min | 1 | $^{5E}$U, 60 min |
| **5** | Nanopore | K562 | $^{5E}$U 60 min | 2 | $^{5E}$U, 60 min |
| **6** | Nanopore | K562 | $^{5E}$U 60 min | 3 | $^{5E}$U, 60 min |
| **7** | Nanopore | K562 | $^{5E}$U 60 min | 4 | $^{5E}$U, 60 min |
| **8** | Nanopore | K562 | Control | 1 | - |
| **9** | Nanopore | K562 | Control | 2 | - |
| **10** | Nanopore | K562 | Control | 3 | - |
| **11** | Nanopore | K562 | $^{5E}$U 24 h | 1 | $^{5E}$U, 24 h (8 h intervals) |
| **12** | Nanopore | K562 | $^{5E}$U 24 h | 2 | $^{5E}$U, 24 h (8 h intervals) |
| **13** | Nanopore | K562 | $^{5E}$U 24 h | 3 | $^{5E}$U, 24 h (8 h intervals) |
| **14** | Nanopore | K562 | $^{5E}$U 60 min HS | 1 | $^{5E}$U, 60 min, 42 °C, 65 min. |
| **15** | Nanopore | K562 | $^{5E}$U 60 min HS | 2 | $^{5E}$U, 60 min, 42 °C, 65 min. |
| **16** | Nanopore | K562 | $^{5E}$U 60 min HS | 3 | $^{5E}$U, 60 min, 42 °C, 65 min. |
| **17** | RNA-seq | K562 | Ctrl | 1 | - |
| **18** | RNA-seq | K562 | Ctrl | 2 | - |

**Supplementary Table 2. Sequencing statistics of 16 direct RNA 'long-read' nanopore sequencing libraries generated in this study.** All libraries were sequenced on a MinION Mk1B (MIN-101B) for 48 h, unless reads sequenced per second stagnated dramatically. Numbers refer to experimental conditions listed in **Supplementary table 1**.

| **No.** | **Condition name** | **Read counts** | | | **Sequencing time** |
|---|---|---|---|---|---|
| | | **Measured** | **Base-called** | **Aligned** | |
| **1** | Synthetic control | 800.475 | 464.568 (58%) | 175.389 (22%) | 20 h |
| **2** | $^{5Br}$U, $^{4S}$U, $^{6S}$G | 42.131 | 30.571 (73%) | 8.652 (21%) | 4 h |
| **3** | $^{5E}$U, $^{5I}$U | 138.119 | 129.516 (94%) | 27.348 (20%) | 21 h |
| **4** | $^{5E}$U 60 min | 287.528 | 251.468 (87%) | 105.841 (37%) | 48 h |
| **5** | $^{5E}$U 60 min | 430.352 | 376.720 (88%) | 139.965 (33%) | 48 h |
| **6** | $^{5E}$U 60 min | 169.197 | 143.426 (85%) | 33.678 (20%) | 48 h |
| **7** | $^{5E}$U 60 min | 624.550 | 622.665 (100%) | 447.186 (72%) | 48 h |
| **8** | Control | 243.032 | 217.405 (89%) | 118.115 (49%) | 48 h |
| **9** | Control | 288.320 | 264.648 (92%) | 141.924 (49%) | 48 h |
| **10** | Control | 280.010 | 244.878 (87%) | 116.903 (42%) | 48 h |
| **11** | $^{5E}$U 24 h | 177.088 | 131.720 (74%) | 28.758 (16%) | 48 h |
| **12** | $^{5E}$U 24 h | 804.756 | 538.046 (67%) | 156.643 (19%) | 48 h |
| **13** | $^{5E}$U 24 h | 106.173 | 80.914 (76%) | 15.098 (14%) | 48 h |
| **14** | $^{5E}$U 60 min HS | 550.323 | 450.982 (82%) | 220.304 (40%) | 48 h |
| **15** | $^{5E}$U 60 min HS | 260.283 | 227.131 (87%) | 99.497 (38%) | 48 h |
| **16** | $^{5E}$U 60 min HS | 204.950 | 188.327 (92%) | 86.284 (42%) | 48 h |

**Supplementary Table 3. List of synthetic RNAs used in this study.** Synthetic RNAs are derived from selected synthetic sequences of the ERCC Spike-in Mix (**Methods**). Synthetic RNAs are polyadenylated after IVT.

| No. | Name | Length (nt) | Derived from | %GC | %U |
|---|---|---|---|---|---|
| 1 | RNA spike-in 2 | 982 | Synthetic, ERCC-00043 | 34 | 30 |
| | 5'-<br>GGGTGCTTTAACAAGAGGAAATTGTGTTTTTGCCAATTTAAGACCTAATTTAATAGTTAAACCATTAACCTTAGT<br>TGTTCCAAGGCATAATATAGAGAGTGAGATACAGGATGAGCTATTTCAGGGAGTTATTCAGTATGCAGTTGCCA<br>AGGCAGTTGCTGATTTAGATTTAGATGAAGATTTAAAGGTTGTTGTCTCTGTTAATGTCCCAGAGGTTCCAATAA<br>CCAATTTAAATAAAAGAAAACTCTTCCAATACTTCTATGCCTCAGCAAAGTTAGCTATAAACAGAGCTTTAAAT<br>GAATATCCTTCAAAAGAGAAGGTAAAGAAAGAGAAATATAGAGCTTTGCATCCATTAGTTGGATTTAGGGATGT<br>TAGATTGGAGTATCCTCCATATCTACAAATTGCTTTGGATGTCCCAACTATGGAGAATTTGGAATTTTTGTTACA<br>AACAATTCCAAATAGCGACCACATCATCTTAGAGGCTGGAACACCACTAATTAAAAAGTTTGGTTTAGAGGTTA<br>TTGAAATAATGAGAGAATATTTTGATGGCTTTATTGTTGCTGATTTAAAAACCTTAGACACTGGAAGGGTTGAG<br>GTAAGATTGGCATTTGAAGCAACAGCTAATGCAGTGGCAATAAGTGGAGTAGCACCAAAATCAACAATAATTA<br>AAGCTATCCACGAATGTCAAAAATGTGGTTTAATCAGCTATTTGGATATGATGAACGTCTCTGAACCTCAAAAA<br>TTATATGATTCATTAAAATTAAAGCCAGATGTTGTTATCTTGCATAGAGGGATTGATGAGGAGACATTTGGAATT<br>AAAAAGGAATGGAAATTTAAGGAAAACTGCTTATTAGCAATTGCTGGAGGAGTTGGTGTGGAGAATGTTGAAG<br>AGCTTTTAAAAGAATATCAAATATTAATCGTTGGTAGAGCAATTACAAAATCAAAAGACCCAGGAAGAGTAATT<br>AGGATTTTATAAACAAGATGG-3' | | | | |
| 2 | RNA spike-in 4 | 1011 | Synthetic, ERCC-00136 | 43 | 26 |
| | 5'-<br>GGGTTTCGACGTTTTGAAGGAGGGTTTTAAGTAATGATCGAGATTGAAAAACCAAAAATCGAAACGGTTGAAAT<br>CAGCGACGATGCCGAATTTGGTAAGTTTGTCGTAGAGCCACTTGAGCGTGGATATGGTACAACTCTGGGTAACT<br>CCTTACGTCGTATCCTCTTATCCTCACTCCCTGGTGCCGCTGTAACATCAATCCAGATAGATGGTGTACTGCACG<br>AATTCTCGACAATTGAAGGCGTTGTGGAAGATGTTACAACGATTATCTTACACATTAAAAAGCTTGCATTGAAA<br>ATCTACTCTGATGAAGAGAAGACGCTAGAAATTGATGTACAGGGTGAAGGAACTGTAACGGCAGCTGATATTA<br>CACACGATAGTGATGTAGAGATCTTAAATCCTGATCTTCATATCGCGACTCTTGGTGAGAATGCGAGTTTCCGAG<br>TTCGCCTTACTGCTCAAAGAGGACGTGGGTATACGCCTGCTGACGCAAACAAGAGAGGCGATCAGCCAATCGGC<br>GTGATTCCGATCGATTCTATCTATACGCCAGTTTCCCGTGTATCTTATCAGGTAGAGAACACTCGTGTAGGCCAA<br>GTTGCAAACTATGATAAACTTACACTTGATGTTTGGACTGATGGAAGCACTGGACCGAAAGAAGCAATTGCGCT<br>TGGTTCAAAGATTTTAACTGAACACCTTAATATATTCGCTGGTTTAACTGACGAAGCTCAACATGCTGAAATCAT<br>GGTTGAAGAAGAAGAAGATCAAAAAGAGAAAGTTCTTGAAATGACAATTGAAGAATTGGATCTTTCTGTTCGTT<br>CTTACAACTGCTTAAAGCGTGCGGGTATTAACACGGTTCAAGAGCTTGCGAACAAGACGGAAGAAGATATGAT<br>GAAAGTTCGAAATCTAGGACGCAAATCACTTGAAGAAGTGAAAGCGAGACTAGAAGAACTTGGACTCGGACTT<br>CGCAAAGACGATTGACTAGTTTCCCTTGTGAACTAGGATTTTCCCGGGTAC-3' | | | | |
| 3 | RNA spike-in 5 | 1012 | Synthetic, ERCC-00145 | 46 | 26 |
| | 5'-<br>GGGACTGTCCTTTCATCCATAAGCGGAGAAAGAGGGAATGACATTGTTCTTACACGGCACAAGCAGACAAAATC<br>AACATGGTCATTTAGAAATCGGAGGTGTGGATGCTCTCTATTTAGCGGAGAAATATGGTACACCTCTTTACGTAT<br>ATGATGTGGCTTTAATACGTGAGCGTGCTAAAAGCTTTAAGCAGGCGTTTATTTCTGCAGGGCTGAAAGCACAG<br>GTGGCATATGCGAGCAAAGCATTCTCATCAGTCGCAATGATTCAGCTCGCTGAGGAAGAGGGACTTTCTTTAGA<br>TGTCGTATCCGGAGGAGAGCTATATACGGCTGTTGCAGCAGGCTTTCCGGCAGAACGCATCCACTTTCATGGAA<br>ACAATAAGAGCAGGGAAGAACTGCGGATGGCGCTTGAGCACCGCATCGGCTGCATTGTGGTGGATAATTTCTAT<br>GAAATCGCGCTTCTTGAAGACCTATGTAAAGAAACGGGTCACTCCATCGATGTTCTTCTTCGGATCACGCCCGG<br>AGTAGAAGCGCATACGCATGACTACATTACAACGGGCCAGGAAGATTCAAAGTTTGGTTTCGATCTTCATAACG<br>GACAAACTGAACGGGCCATTGAACAAGTATTACAATCGGAACACATTCAGCTGCTGGGTGTCCATTGCCATATC<br>GGCTCGCAAATCTTTGATACGGCCGGTTTTGTGTTAGCAGCGGAAAAAATCTTCAAAAAACTAGACGAATGGAG<br>AGATTCATATTCATTTGTATCCAAGGTGCTGAATCTTGGAGGAGGTTTCGGCATTCGTTATACGGAAGATGATGA<br>ACCGCTTCATGCCACTGAATACGTTGAAAAAATTATCGAAGCTGTGAAAGAAAATGCTTCCCGTTACGGTTTTG<br>ACATTCCGGAAATTTGGATCGAACCGGGCCGTTCTCTCGTGGGAGACGCAGGCACAACTCTTTATACGGTTGGC<br>TCTCAAAAAGAAGTGGATAAGCTGTACAATCGTTTCATCATTCGGCGTGCG-3' | | | | |
| 4 | RNA spike-in 8 | 1076 | Synthetic, ERCC-00092 | 52 | 27 |

| | | | | | |
|---|---|---|---|---|---|
| | 5'-<br>GGGGATGTCCTTGGACGGGGTGGCGCAGTATTACTGCAAGAGAGCGGACAGATTAGTGTGTTGGAGCCGACAC<br>ATCAAAGGTTCGTCCGGGGACCGATCTGCAGCCTACGGGACATTTATCCGTAAAAGCATGGCGCTGTTTCGTAC<br>TTATCGGAGGCCAGGTATCGTCGCGGCGAGTCTCCCCGACGACGGAGATGGGCGTTACTATCTGGGCCGTCTCG<br>TACTCTGTTACTTGGCACAGATGCGAGCCCTCGTAATGTGCATCAGCTAAGGGCGATATTATAATGCGACGTTTG<br>TACGGATTCGTTACTAACGTGTTGGACGCTAGTGGAATATGTGTCGTTGGTTAGCCTACCCATGGCTTTCGCGGC<br>GACACATGCTTAGACTCTTTCAAAACTTCGGTGAAGTTCACTCAAGCCGCGGAGCGCCGTCGTAATTCACTAGG<br>GATGGCGGTACCCGTGCCCGTCCGATTCGTAGCAACCTGCATCACGATTTTGTCTTCGGGCGACTTATCAGATAC<br>GGTAATGTAAATACCTGGCATTTGGGCACTTCTTGCGTTTAAGCGGGAAAGATCGCGAGGGCCCGCTATTTGCG<br>ATACTTCCCATGTCGGTGCCGTCGCCTCTATGTACTCGGAGACGTTAATGCAGAGGCTAAGGACAATTTACCATG<br>ACTCGGTAATCCGTTCGTCAAGCAGGTAGCTCGAGTCTCCCCACGGACACGTAGTGGGTTTGTAACGATCGATA<br>CCGAGTCTTTTTGTCTAGTAGAACCAACCAACCATTAAGGAGTTCACTAGCACATCTTTGCGACCCGATCGTCCG<br>TGTGTCGCGTAATACTTTTGTTATGACGAGACATACGCTCAAGCCCTGGGTAGCTAGTCGCGGAGGCACGTTAC<br>CGCGCACAACCCCTATTCGTTTACATGTACATCGCATCTGAGGTAGTACACTTCCGGCGTACGTGAGTATTTGCG<br>CGTAATAAGCGCGTGTTTAGCTGATCCCCTCTCGTATCGAGGTTAAGGCAGATTAGTGCCCAGTAATTGCGTTTT<br>TTTGTCGTTGTCGCAGAACGCGATTTGCTCCGAAAGC-3' | | | | |
| 5 | RNA spike-in 9 | 1034 | Synthetic, ERCC-00002 | 53 | 25 |
| | 5'-<br>GGGCCAGATTACTTCCATTTCCGCCCAAGCTGCTCACAGTATACGGGCGTCGGCATCCAGACCGTCGGCTGATC<br>GTGGTTTTACTAGGCTAGACTAGCGTACGAGCACTATGGTCAGTAATTCCTGGAGGAATAGGTACCAAGAAAAA<br>AACGAACCTTTGGGTTCCAGAGCTGTACGGTCGCACTGAACTCGGATAGGTCTCAGAAAAACGAAATATAGGCT<br>TACGGTAGGTCCGAATGGCACAAAGCTTGTTCCGTTAGCTGGCATAAGATTCCATGCCTAGATGTGATACACGT<br>TTCTGGAAACTGCCTCGTCATGCGACTGTTCCCCGGGGTCAGGGCCGCTGGTATTTGCTGTAAAGAGGGGCGTT<br>GAGTCCGTCCGACTTCACTGCCCCCTTTCAGCCTTTTGGGTCCTGTATCCCAATTCTCAGAGGTCCCGCCGTACG<br>CTGAGGACCACCTGAAACGGGCATCGTCGCTCTTCGTTGTTCGTCGACTTCTAGTGTGGAGACGAATTGCCAGA<br>ATTATTAACTGCGCAGTTAGGGCAGCGTCTGAGGAAGTTTGCTGCGGTTTCGCCTTGACCGCGGGAAGGAGACA<br>TAACGATAGCGACTCTGTCTCAGGGGATCTGCATATGTTTGCAGCATACTTTAGGTGGGCCTTGGCTTCCTTCCG<br>CAGTCAAAACCGCGCAATTATCCCCGTCCTGATTTACTGGACTCGCAACGTGGGTCCATCAGTTGTCCGTATACC<br>AAGACGTCTAAGGGCGGTGTACACCCTTTTGAGCAATGATTGCACAACCTGCGATCACCTTATACAGAATTATC<br>AATCAAGCTCCCCGAGGAGCGGACTTGTAAGGACCGCCGCTTTCGCTCGGGTCTGCGGGTTATAGCTTTTCAGTC<br>TCGACGGGCTAGCACACATCTGGTTGACTAGGCGCATAGTCGCCATTCACAGATTTGCTCGGCAATCAGTACTG<br>GTAGGCGTTAGACCCCGTGACTCGTGGCTGAACGGCCGTACAACTCGACAGCCGGTGCTTGCGTTTTACCC-3' | | | | |
| 6 | RNA spike-in 12 | 947 | Synthetic, ERCC-00170 | 35 | 31 |
| | 5'-<br>GGGGCACAAGTTGCTGAAGTTGCGAGAGGGGCGATAAGTGAGGCAGACAGGCATAATATAAGAGGGGAGAGA<br>ATTAGCGTAGATACTCTTCCAATAGTTGGTGAAGAAAATTTATATGAGGCTGTTAAAGCTGTAGCAACTCTTCCA<br>CGAGTAGGAATTTTAGTTTTAGCTGGCTCTTTAATGGGAGGGAAGATAACTGAAGCAGTTAAAGAATTAAAGGA<br>AAAGACTGGCATTCCCGTGATAAGCTTAAAGATGTTTGGCTCTGTTCCTAAGGTTGCTGATTTGGTTGTTGGAGA<br>CCCATTGCAGGCAGGGGTTTTAGCTGTTATGGCTATTGCTGAAACAGCAAAATTTGATATAAATAAGGTTAAAG<br>GTAGGGTGCTATAAAGATAATTTAATAATTTTTGATGAAACCGAAGCGTTAGCTTTGGGTTATGAAACTCCATG<br>ATTTTCATTTAATTTTTTCCTATTAATTTTCTCCTAAAAAGTTTCTTTAACATAAATAAGGTTAAAGGGAGAGCTC<br>TATGATTGTCTTCAAAAATACAAAGATTATTGATGTATATACTGGAGAGGTTGTTAAAGGAAATGTTGCAGTTG<br>AGAGGGATAAAATATCCTTTGTGGATTTAAATGATGAAATTGATAAGATAATTGAAAAAATAAAGGAGGATGTT<br>AAAGTTATTGACTTAAAAGGAAAATATTTATCTCCAACATTTATAGATGGGCATATACATATAGAATCTTCCCAT<br>CTCATCCCATCAGAGTTTGAGAAATTTGTATTAAAAAGCGGAGTTAGCAAAGTAGTTATAGACCCGCATGAAAT<br>AGCAAATATTGCTGGAAAAGAAGGAATTTTGTTTATGTTGAATGATGCCAAAATTTTAGATGTCTATGTTATGCT<br>TCCTTCCTGTGTTCCAGCTACAAACTTAGAAACAAGTGGAGCTGAGATTACAGCAGA-3' | | | | |

**Supplementary Table 4. List of previously published datasets used in this study.**

| No. | Data | Cell line | Used for | Reference | Available at |
|---|---|---|---|---|---|
| 1 | GRO-cap | K562 | Full length isoform assessment | Core et al. 2014 [3] | NCBI Gene Expression Omnibus (GSE60456) |

Data accession links

1: https://www.ncbi.nlm.nih.gov/geo/query/acc.cgi?acc=GSE60456
